# FluoMALDI microscopy: matrix co-crystallization simultaneously enhances fluorescence and MALDI imaging

**DOI:** 10.1101/2023.05.27.542340

**Authors:** Ethan Yang, Xinyi Elaine Shen, Hoku West-Foyle, Dalton R. Brown, Cole C. Johnson, Jeong Hee Kim, LaToya Ann Roker, Caitlin M. Tressler, Ishan Barman, Scot C. Kuo, Kristine Glunde

## Abstract

We report that co-crystallization of fluorophores with matrix-assisted laser desorption/ionization (MALDI) imaging matrices significantly enhances fluorophore brightness up to 79-fold, enabling the amplification of innate tissue autofluorescence. This discovery facilitates FluoMALDI, the imaging of the same biological sample by both fluorescence microscopy and MALDI imaging. Our approach combines the high spatial resolution and specific labeling capabilities of fluorescence microscopy with the inherently multiplexed, versatile imaging capabilities of MALDI imaging. This new paradigm eliminates the notion that MALDI matrices obscure and obstruct optical microscopy approaches, allowing to image the exact same cells in tissues, free of any physical changes between fluorescence and MALDI imaging, which minimizes data registration processes. Matrix-fluorophore co-crystallization also facilitates applications with insufficient fluorescence brightness. We showcase the capabilities of FluoMALDI imaging with endogenous and exogenous fluorophores and autofluorescence-based FluoMALDI of brain and kidney tissue sections. FluoMALDI will advance structural-functional microscopic imaging in cell biology, biomedicine, and pathology.

## INTRODUCTION

Mass spectrometry imaging (MSI) is a powerful technology which uses an intrinsic molecular parameter, the molecular mass, to directly identify and visualize the distribution of a wide variety of biomolecules in tissue sections, cells, and other complex environments. MSI achieves highly multiplexed imaging without requiring specific labels or knowledge of the tissue’s biochemical composition and has consequently evolved into an important molecular discovery tool^1^. Owing to the wide mass range, soft ionization to generate molecular ions, and advances in speed and spatial resolution, matrix-assisted laser desorption/ionization (MALDI) imaging is currently the most frequently used MSI tool in biomedical and clinical research with wide-ranging applications^1^. In MALDI imaging, a tissue section is raster-scanned with a laser in a grid-like fashion to generate a mass spectrum of the molecular composition of each pixel within the scanned region^1, 2^. Samples prepared for MALDI imaging are coated with a thin layer of matrix crystals, which co-crystallize with proximal analyte molecules^3-6^. During MALDI imaging, matrix-analyte co-crystals readily absorb ultraviolet light from a focused laser beam and thereby enhance desorption and ionization of the analyte molecules from the tissue section surface^3-6^. MALDI imaging achieves spatial resolutions of 5 µm on commercial instruments^7^, and 1.4 µm on home-built instrumentation^8^. Transmission-mode MSI with MALDI-2 post-ionization for signal enhancement has achieved subcellular resolution of 600 nm^9^. Currently, clinical applications of MALDI imaging are rapidly developing because of their capability to integrate with pathology as well as their ability to provide molecular biomarker profiles for diagnosis, prognosis, and response to therapy in several diseases^2, 10^. As of today, it is possible to acquire MALDI images of metabolites^11^ and metabolic activity^12^, lipids^13-15^, drugs and drug metabolites^16^, amino acids and neurotransmitters^7, 17^, native and tryptic peptides following on-tissue tryptic digest^18^, and other on-tissue enzyme-digested biomolecules including N-glycans^19^ and collagens^20^.

Optical microscopy techniques such as colorimetric histology staining^21^ and fluorescence microscopy^22^ have been widely utilized for clinical and research applications. Moreover, optical microscopy techniques have enabled large-scale tissue atlas building projects such as the Allen brain atlas^23^, the human protein atlas^24^, and the human kidney atlas^25^, with others currently being constructed^26^. These atlases contain vast amounts of publicly available anatomical and morphological data. Current MALDI imaging workflows frequently utilize and refer to this existing knowledge by acquisition of optical images to help navigate and interpret the corresponding MALDI imaging data^27-30^. Combining autofluorescence approaches^27, 28^ with MALDI imaging is particularly promising, as it reveals detailed anatomical, morphological, and cellular structural information without the need for any staining or other sample preparation protocols^11,28,31^.

In current workflows, the autofluorescence microscopy is performed prior to MALDI matrix coating and MALDI imaging, due to the belief that the matrix obscures and obstructs optical microscopy approaches. To achieve anatomical and morphological referencing of MALDI imaging with corresponding optical microscopy data, it is critical to correctly co-register the two datasets. Many current methods still rely on manually aligning datasets based on intensity patterns and fiducial markers^32, 33^, introducing bias and inaccuracies. Recent developments have improved the accuracy and speed of the registration process between datasets through computational approaches, using image fusion^30^ and registering MSI pixels to optically detected laser ablation patterns^28^. However, such multimodal image registration and fusion methods are limited by time-consuming computation and require specialized, often home-built software.

In this study, we present the FluoMALDI workflow, which is made possible by our paradigm-shifting discovery that co-crystallization of MALDI matrices with fluorophores significantly enhances their brightness. The FluoMALDI pipeline integrates fluorescence and MALDI imaging while providing enhanced fluorescence intensity and increased molecular information from the same tissue section. In the FluoMALDI pipeline, we perform matrix coating before acquisition of fluorescence images, which is then followed by MALDI imaging, overcoming many technical issues and limitations with current fluorescence and MALDI imaging workflows. FluoMALDI produces (1) great enhancement in fluorescence intensities, (2) allows the full experiment to be performed on the same tissue section, and (3) achieves simple alignment between data sets through linear registration. This novel pipeline can acquire highly multiplexed information on biological samples through autofluorescence-guided MALDI imaging. FluoMALDI only requires one tissue section and one sample preparation for the entire workflow, thereby circumventing technical challenges arising from staining and complex data registration. The discovery of FluoMALDI also creates new paradigms for combined instrumentation for fluorescence and MALDI imaging, which could further reduce the time required for FluoMALDI imaging experiments in the future, opening up new avenues for direct morphology-guided MALDI imaging acquisition.

## RESULTS

### FluoMALDI Imaging Pipeline

The FluoMALDI multimodal imaging pipeline is depicted in **Fig. 1**. First, biological samples are cryosectioned and thaw-mounted onto indium-tin-oxide (ITO) microscopy slides. Next, a MALDI matrix of choice is deposited on samples to achieve a thin layer of matrix coating at the desired thickness. Fluorescence microscopy is then performed at the desired wavelength(s) with either a widefield or confocal microscope. MALDI imaging data is then acquired on the same slide with the desired parameters.

**Figure 1.**
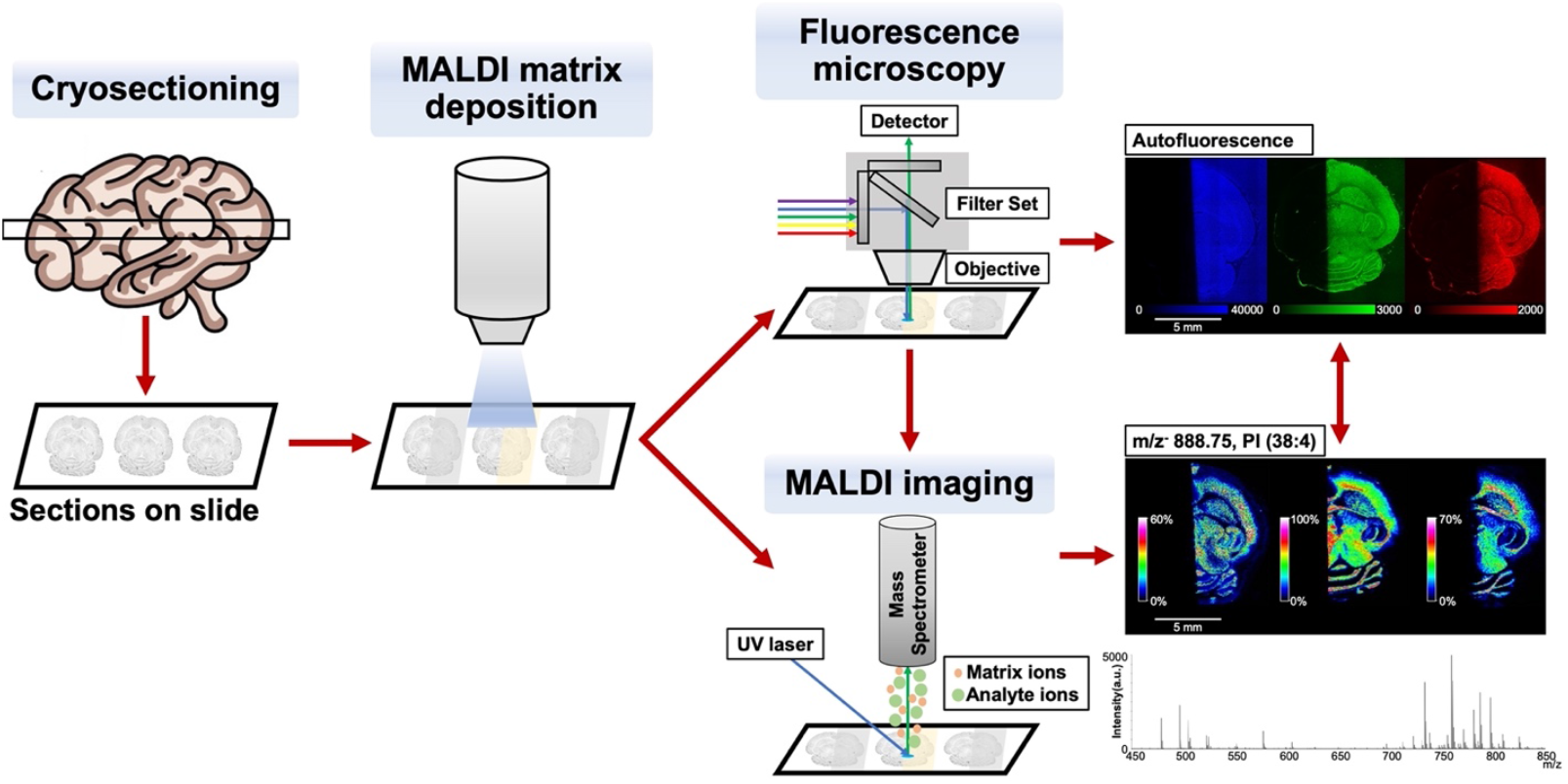
Experimental workflow for FluoMALDI microscopy. A fresh-frozen tissue, in this example mouse brain, is cryo-sectioned and thaw-mounted onto a ITO microscopy slide. This is followed by MALDI matrix deposition, i.e., the tissue sections are sprayed with the MALDI matrix of choice, in this case on one half of the brain only. The matrix-coated tissue sections are imaged with fluorescence microscopy, and then subjected to MALDI MSI. This workflow enables the seamless integration of (auto)fluorescence and MALDI imaging data, while simultaneously enhancing fluorescence and MALDI MSI signals.

The alignment of fluorescence microscopy and MALDI imaging data and quantification of fluorescence and MALDI MS signals is performed by simple linear registration using ImageJ^34^ or other image analysis software. Quantification of fluorescence and MALDI MSI signals can proceed on the registered dataset.

### MALDI Matrices Enhance Fluorescence Signal Intensities of Fluorophores

To demonstrate and quantitatively measure the enhanced fluorescence intensities from MALDI matrix coating of fluorescent samples in FluoMALDI, we first performed the workflow shown in Fig. 1 with pink Sharpie® permanent marker drawings of “J”‘s on an ITO coated microscopy slide. Pink Sharpie® marker was used because it contains the fluorophore Rhodamine B and/or 6G (Rhodamine B/6G), both of which are isomeric and isobaric when detected by MALDI imaging (**Fig. S1**)^35, 36^. Matrix coatings were deposited with a robotic pneumatic sprayer in all experiments in our study. We observed significant fluorescence signal enhancements for all pink Sharpie® J’s that were coated with any of six common MALDI matrices including 1,5-diaminonapthalene (DAN), 9-aminoacridine (9AA), 2,5-dihydroxybenzoic acid (DHB), sinapic acid (SA), norharmane (nH), and α-cyano-4-hydroxycinnamic acid (CHCA), as measured with a red fluorescence filter set (TRITC, 543/22 nm excitation, 593/40 nm emission) and shown in **Fig. 2**. Representative images of the Sharpie® J marks with each matrix coating compared to uncoated and solvent-sprayed controls are shown in **Fig. 2A**. Solvent spraying alone did not have any effect on fluorescence signal intensity, while all tested MALDI matrix coatings caused significant enhancements in red fluorescence signal, with the greatest enhancement observed for nH^15^ (37-fold, n=3, **Fig. 2B-C**) and CHCA^37^ (28-fold, n=3, **Fig. 2B-C**).

**Figure 2.**
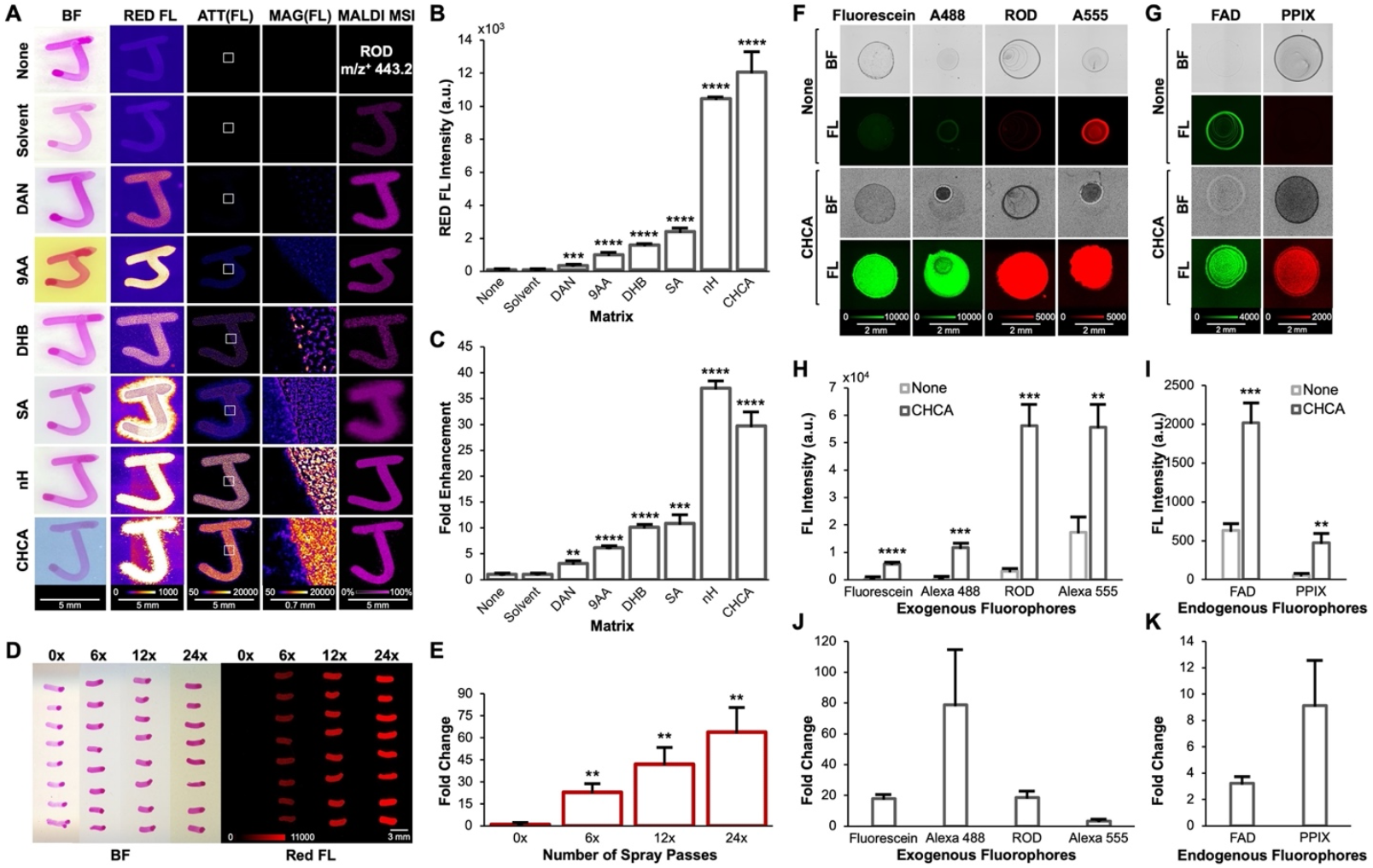
MALDI matrix coating of fluorophores results in fluorescence intensity enhancement. (**A**) Columns from left to right show optical brightfield scans, red fluorescence images, attenuated red fluorescence images, magnified red fluorescence images from regions of interest (ROIs) within white squares, and MALDI images at m/z 443.2 Da corresponding to the [M]^+^ ion of Rhodamine B/6G (see **Fig. S1** for tandem MS data). Rows show uncoated controls and coating with various MALDI matrices at 1.6 µg/mm^2^ density. Fluorescence and MALDI images are shown on the same intensity scale per column. (**B**) Quantification of fluorescence intensities using various matrices on Rhodamine B/6G from (A), n=3. (**C**) Fold enhancement of red fluorescence from Rhodamine B from each matrix coating, normalized to uncoated control (none), n=3. (**D**) Enhancement of pink Sharpie® lines coated with increasing pass numbers of CHCA spray, ranging from 0 passes (0x, 0 µg/mm^2^) to 24 passes (24x, 1.6 µg/mm^2^). (**E**) Corresponding fold fluorescence enhancement of CHCA matrix coating at varying densities (numbers of spray passes) normalized to uncoated samples (0x). (**F**) Red or green fluorescence enhancements of exogenous fluorophores including Fluorescein, Alexa (A) 488, Rhodamine B (ROD), and A555, as coated with CHCA versus uncoated controls (none). (**G**) Fluorescence enhancements of endogenous fluorophores including flavin adenine dinucleotide (FAD) and protoporphyrin IX (PPIX), as coated with CHCA versus uncoated controls (none). (**H**) Quantification of fluorescence intensities from exogenous fluorophores shown in (F), n=3. (**I**) Quantification of fluorescence intensities from endogenous fluorophores shown in (G), n=3. (**J**) Fold enhancement for each exogenous fluorophore shown in F, H. (**K**) Fold enhancement for each endogenous fluorophore shown in G, I. All quantitative data are shown as mean values ± standard error of three independent experiments. ** p<0.01, *** p<0.001, **** p<0.0001. Abbreviations: ATT, attenuated; BF, brightfield; FL, fluorescence; MAG, magnified.

Next, we sought to determine if the sprayed matrix density (i.e., number of sprayed matrix layers) had any correlation with fluorescence signal intensity. We drew horizontal lines of pink Sharpie® permanent marker onto microscopy slides, which were then sprayed with CHCA at different densities ranging from 0 µg/mm^2^ (i.e., 0 layers or 0x) to 1.6 µg/mm^2^ (i.e., 24 layers or 24x), followed by widefield fluorescence microscopy using the red fluorescence filter set (**Fig. 2D**). Quantification of this data demonstrated that the resulting fluorescence signal intensity increased linearly with matrix density (**Fig. 2E**). Typical CHCA matrix densities used for MALDI imaging are 0.5 µg/mm^2^ (i.e., 8 layers or 8x).

Next, we tested if the observed fluorescence enhancement phenomenon would consistently occur with other fluorophores. CHCA coating (1.6 µg/mm^2^) was applied to four commonly used exogenous fluorophores spotted on microscopy slides at a concentration of 0.2 mg/mL each (**Fig. 2F**). Representative images of Fluorescein and Alexa Fluor 488 (A488) were acquired using a green fluorescence filter set (GFP, 472/30 nm excitation, 520/35 nm emission), while Rhodamine B and Alexa Flour 555 (A555) were imaged using a red fluorescence filter set. All CHCA-coated fluorophore spots displayed significantly increased fluorescence signal intensity compared to uncoated control spots, with Rhodamine B and A555 showing the highest signal intensities (**Fig. 2H**). The fold enhancement ranged between 3-fold for A555 to 79-fold for A488.

We also investigated the effect of matrix coating on endogenous fluorophores, including flavin adenine dinucleotide (FAD) and protoporphyrin IX (PPIX), both of which are ubiquitous in cells and significantly contribute to innate tissue autofluorescence signals^38^. CHCA was deposited onto FAD spots and imaged using the green filter set, as well as onto PPIX spots which were imaged using the red filter set (**Fig. 2G**). Similar to the tested exogenous fluorophores, matrix coating also significantly increased the brightness of endogenous fluorophores, with 3-fold enhancement for FAD and 9-fold enhancement for PPIX (**Fig. 2I, 2K**). We tested additional exogenous (Fluorescein, Hoechst, A350) and endogenous (NADH, NADPH) fluorophores, acquiring comprehensive fluorescence data of all tested fluorophores using blue (DAPI, 377/50 nm excitation, 447/60 nm emission), green, and red filter sets as reported in **Fig. S2** and **Fig. S3**. Measurements of CHCA- and nH-coated samples in **Figs. S2-S4, S6** revealed significant blue fluorescence from CHCA and nH alone, possibly obscuring blue fluorescing fluorophores (Hoechst, A350, NADH) and blue tissue autofluorescence.

### MALDI Matrices Enhance Autofluorescence Signal Intensities of Tissue Sections

Since endogenous fluorophores displayed significant FluoMALDI enhancements, we next investigated whether MALDI matrix-coating would produce FluoMALDI effects for innate tissue autofluorescence. Starting with transverse (axial) mouse brain tissue sections, autofluorescence was detected without matrix coating in blue, green, and red fluorescence channels, with varying intensities across different anatomical brain regions (**Fig. 3A-B, S4**). The most intense autofluorescence prior to matrix coating was observed in the green channel, as established previously for tissues in general^38^. We sprayed matrix on one half of the brain while the other half was blocked from the spray with a strip of aluminum foil, allowing for direct comparison between matrix-coated and uncoated regions on the same sample. With matrix coating, we observed significant green and red fluorescence intensity enhancements for all matrix-coated halves compared to the matrix-free control halves and uncoated control, without losing anatomic specificity of the respective autofluorescence signal (**Fig. 3A, S4**). Overall, CHCA-, 9AA-, and nH-coated brain sections displayed the most intense fluorescence enhancements, which varied depending on fluorescence channel. Autofluorescence intensities in brain sections coated with nH were the most enhanced in the green channel, while CHCA-coating resulted in the most significant fluorescence enhancement in the red channel (**Fig. 3B**). 9AA caused the strongest signal enhancement in autofluorescence intensity, while nH and CHCA resulted in larger fold enhancements, due to lower background fluorescence from the matrix itself. MALDI imaging in negative ion mode was performed on the same brain sections following fluorescence imaging. Corresponding MALDI imaging data are shown for m/z^-^ 885.75 Da (**Fig. 3A**), which was identified by tandem MS as phosphatidylinositol (PI) (38:4) ([M-H]^-^, see **Fig. S5** or tandem MS data). Comprehensive fluorescence and MALDI imaging data for DAN, DHB, and SA matrix-coated brain tissue sections are reported in **Fig. S5**. Comparable FluoMALDI data were obtained for mouse kidney sections shown in **Fig. S6**, including corresponding tandem MS data in **Fig. S7-S9**.

**Figure 3.**
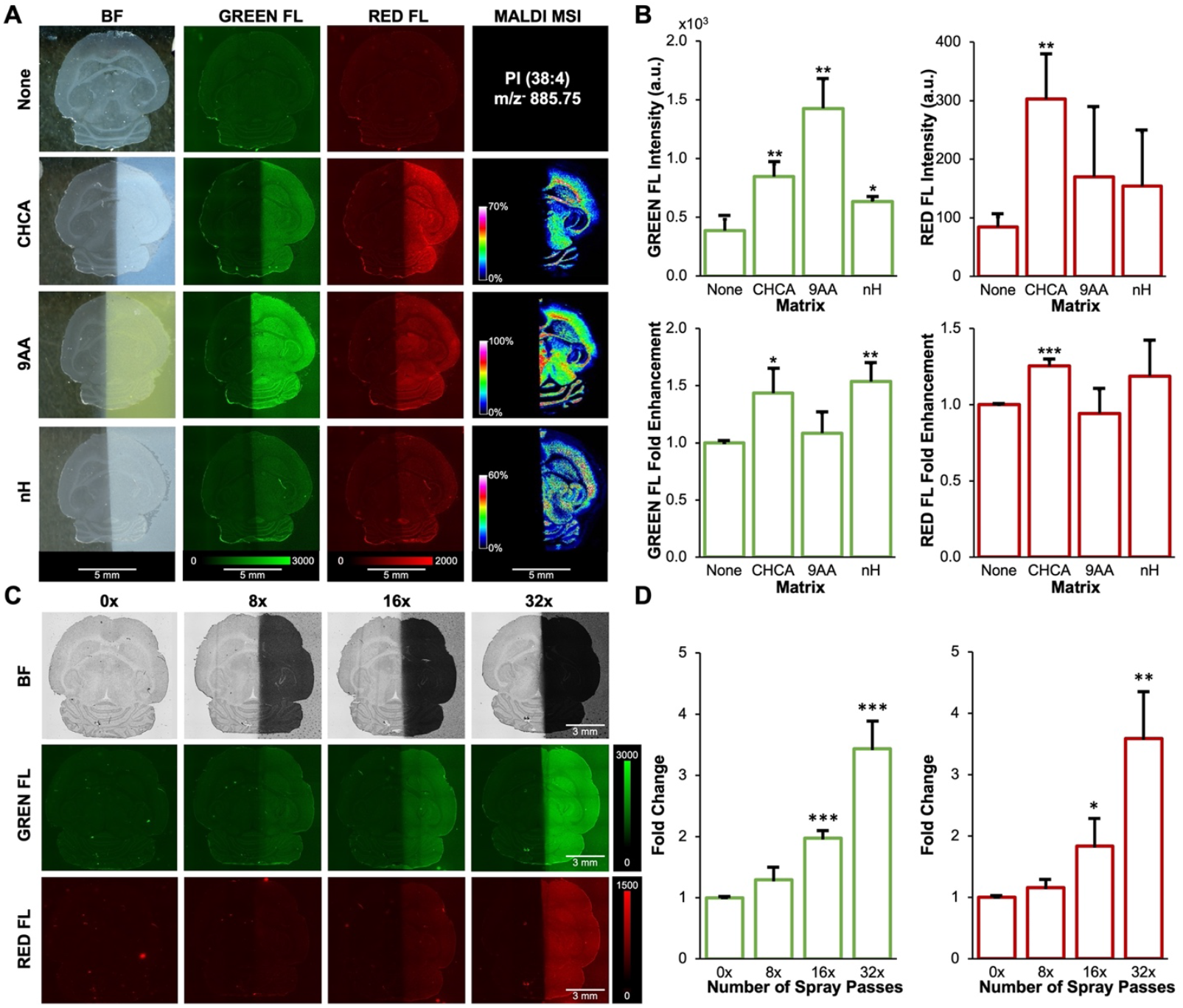
MALDI matrix coating of mouse brain tissue sections increases autofluorescence intensity. (**A**) Imaging results of transverse (axial) mouse brain tissue sections showing, in columns from left to right, brightfield, green fluorescence, red fluorescence, and MALDI imaging in negative ion mode displaying m/z^-^ 885.75 Da, which was identified by tandem MS as phosphatidylinositol (PI) (38:4) ([M-H]^-^, see **Fig. S5** for tandem MS data). Rows show uncoated controls and coating with various MALDI matrices at 1.6 µg/mm^2^ density. Fluorescence and MALDI images are shown on the same intensity scale per column.(B) Quantification of green (left), and red (right) fluorescence intensities and corresponding fold enhancements from (A), n=3.(C)Autofluorescence enhancement of mouse brain sections coated with varying densities of CHCA, ranging from 0 passes (0x, 0 µg/mm^2^) to 32 passes (32x, 2.2 µg/mm^2^). (**D**) Fold change in green (left) and red (right) fluorescence intensity of each matrix density normalized to uncoated controls (0x) corresponding to (C). All quantitative data are shown as mean values ± standard error of three independent experiments. * p<0.05, ** p<0.01, *** p<0.001. Abbreviations: BF, brightfield; FL, fluorescence.

We also investigated the correlation between matrix density and tissue autofluorescence intensity, using the same approach as for pink Sharpie® marks in **Fig. 2**. CHCA was selected as our matrix of choice, as it results in intense fluorescence enhancements in both Sharpie® marks and tissue sections and is widely used for MALDI imaging^37^. With increasing matrix coating density from 0 µg/mm^2^ (i.e., 0 layers or 0x) to 2.2 µg/mm^2^ (i.e., 32 layers or 32x), we observed significant green and red fluorescence intensity enhancements (**Fig. 3C-D**). Tissue integrity and autofluorescent features were preserved with increasing matrix density.

### Co-crystallization of Fluorophores with MALDI Matrices Increases their Fluorescence

We sought to further investigate the observed phenomenon that MALDI matrix co-crystallization with fluorophores enhances the fluorophores’ brightness. To determine if there were structural differences between crystals formed by matrix alone and those formed in the presence of fluorophores, we acquired high (40x) magnification transmitted-light images under crossed polarizers of each matrix, as sprayed on bare glass or over the Rhodamine B/6G Sharpie® marks. As shown in **Figs. 2A** and **2D**, fluorescence is confined to the region of the Sharpie® marks themselves. In the magnified inset in **Fig. 2A**, a bright, granular structure, caused by the fluorophore-matrix co-crystals, is present within the marks, which was not observed in the unsprayed controls. Under higher magnification we clearly observed that the Rhodamine dye is incorporated into the matrix crystals, staining them red (**Fig. 4A**). Furthermore, the structure of the matrix crystals visibly changes when co-crystallized with Rhodamine B/6G, where DAN, 9AA, DHB, and SA show the most noticeable crystal shape differences between matrix-Rhodamine-B/6G co-crystals and pure matrix crystals.

**Figure 4.**
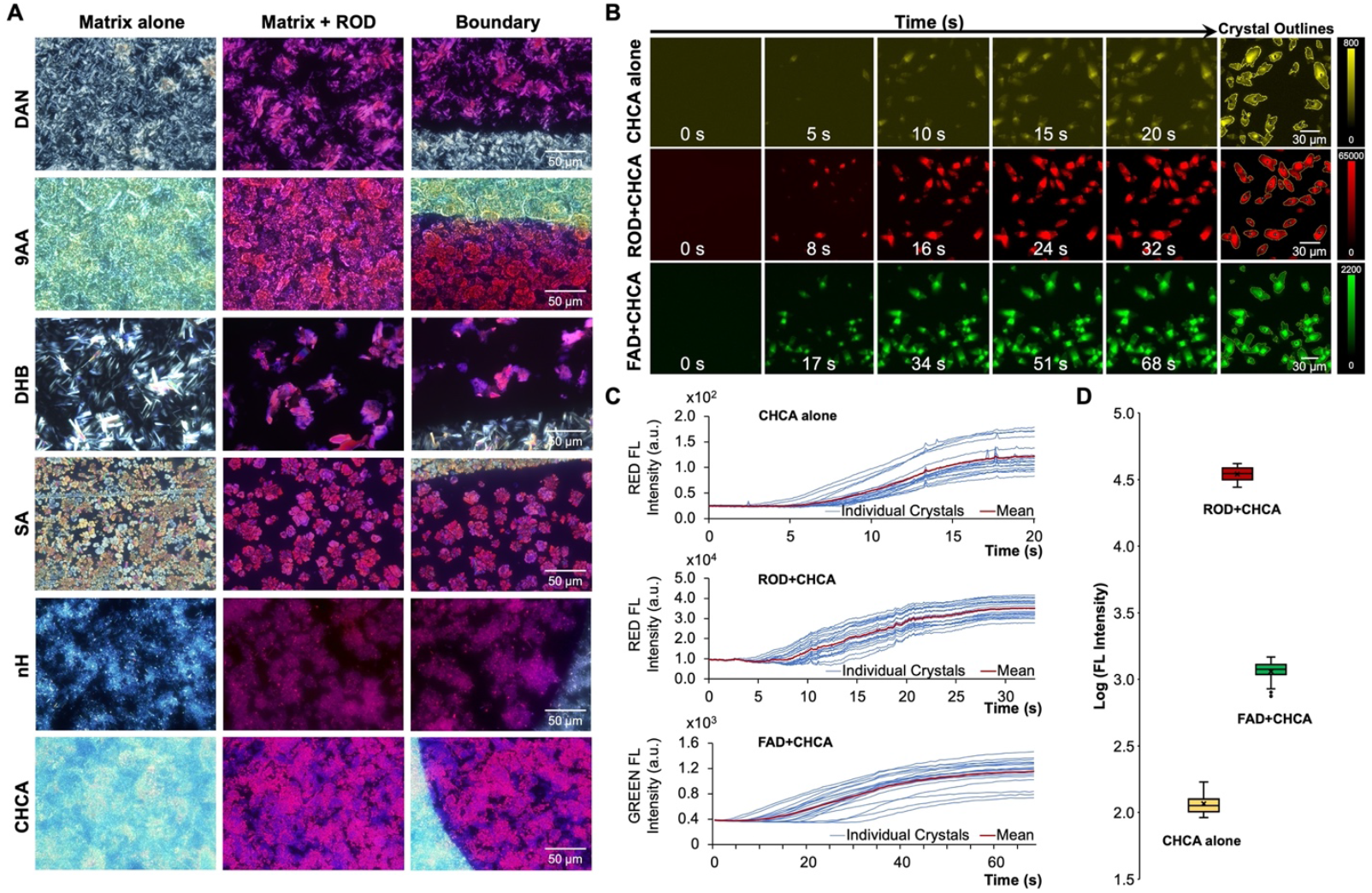
Co-crystallization of fluorophores with matrices. (**A**) From left to right, columns show polarized light microscopy images of matrix crystals, matrix-Rhodamine B/6G (ROD) co-crystals, and the boundary between these two types of crystals acquired with a 40x objective. Rows show coating with various MALDI matrices. (**B**) Representative frames from time-lapse fluorescence microscopy acquisition of crystal formation for CHCA alone (top), co-crystallization of Rhodamine B/6G (ROD) with CHCA (middle), and co-crystallization of FAD with CHCA (bottom). The last column shows the 20 largest crystals selected for quantification for each type of crystallization. (**C**) Time courses of fluorescence intensity quantification of the 20 largest crystals for (co-)crystallization of CHCA alone, ROD+CHCA, and FAD+CHCA. Blue lines represent individual crystals’ fluorescence intensities, red lines are mean crystal fluorescence intensities. The start time of t = 0 s was set to the appearance of the first crystal. A full-length time course including addition of the fluorophore to matrix is shown for FAD+CHCA in **Fig. S10**. (**D**) Fluorescence intensity distributions of the 20 selected crystals each for CHCA alone, ROD+CHCA, and FAD+CHCA shown on a log intensity scale to capture the large differences.

To further examine fluorescence enhancement through co-crystallization, we recapitulated the matrix-fluorophore co-crystallization process, as it would typically occur during matrix spraying, by performing time-lapse fluorescence microscopy experiments where equal volumes of matrix solution and fluorophore solution were mixed, and the co-crystallization process was recorded in real-time (**Supporting Videos V1-V4**).

For CHCA alone in the red fluorescence channel, crystals grew in irregular shapes and quickly, within seconds, gained intensity and leveled at an average intensity of 110 a.u. (**Fig. 4B-D**). For CHCA with Rhodamine B/6G in the red fluorescence channel, co-crystals grew which had the Rhodamine B/6G molecules incorporated, also gaining red fluorescence intensity within seconds, and saturating at an intensity approximately 300-fold brighter than CHCA crystals alone (**Fig. 4B-D**). Finally, with CHCA and FAD in the green fluorescence channel, co-crystals grew within seconds somewhat more gradually, resulting in an average 10-fold increase in green fluorescence intensity when co-crystallization growth was saturated (**Fig. 4B-D**). The fluorescence intensity of the solution surrounding the crystals also decreased concomitant with crystal growth, indicating soluble fluorophore was being depleted from the droplet (see full time course of ROD+CHCA in **Fig. S10** and video **V4**).

### FluoMALDI Pipeline for Enhanced Spectral Imaging using Laser-Scanning Confocal Microscopy

We next expanded the FluoMALDI workflow to acquire spectral imaging data with a confocal laser-scanning fluorescence microscope. As demonstrated in **Fig. 5**, fluorescence intensity enhancements of FluoMALDI clearly enhance spectral imaging acquisition using laser-based confocal fluorescence microscopes. Fluorescence spectral imaging data were acquired on half-coated brain sections using each excitation wavelength available on the microscope in turn, with detection in 9-nm-wide emission bands from 415 nm to 690 nm. Spectral imaging showed significant fluorescence enhancements on the matrix-coated halves compared to the corresponding non-coated halves, with the most striking enhancements in the excitation range of 488 nm and 514 nm (**Fig. 5B**). All other excitation wavelengths produced fluorescence enhancements from matrix-coating as well, but the differences were less remarkable and matrix background was significant at the shorter wavelengths (**Fig. S12**). The increase in fluorescence intensity through matrix-coating allowed for clear FluoMALDI autofluorescence visualization of the hippocampal horn in the matrix-coated halves only, which was not possible in the corresponding uncoated control halves (**Fig. 5B**). MALDI MSI analysis at 5 µm spatial resolution was then performed on the same tissue section, with molecular distributions of metabolites, peptides, and lipids clearly showing the anatomical structures and morphologies of the hippocampal horn region (**Fig. 5D**). We show two representative lipid MS images of LPE (18:2) at m/z^+^ 478.33 Da and PC (O-16:0) at m/z^+^ 504.35 Da (**Fig. 5D)**, which were identified using tandem MS (**Fig. S13-S14**). Expanded mass spectra of two spectrally different regions of interest (ROIs) (white boxes 1, 2 in **Fig. 5D**) are shown in **Fig. 5E**. Spectral imaging of the same ROIs at 488 nm and 514 nm excitation revealed distinct fluorescence emission spectra associated with each ROI. Following MALDI imaging measurement, the sample was hematoxylin-and-eosin (H-E) stained and imaged in color for anatomical-structural confirmation of the analyzed hippocampal horn region (**Fig. 5A, S11)**.

**Figure 5.**
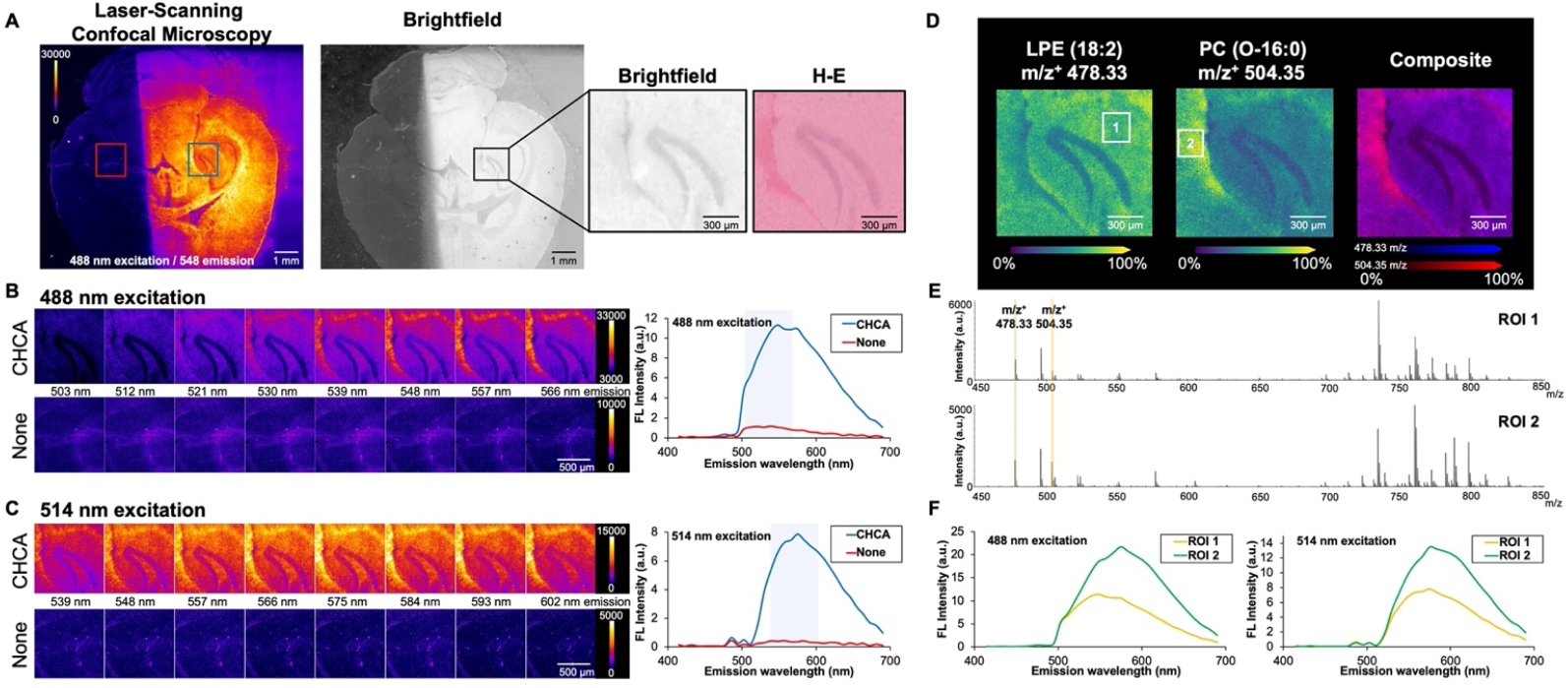
FluoMALDI pipeline using spectral imaging with laser-scanning confocal microscopy and MALDI MSI on mouse brain tissue sections half-coated with CHCA. (**A**) Transverse (axial) half-coated brain section confocally imaged at 488 nm excitation and 548 emission, and corresponding brightfield image with enlarged insets of the hippocampal horn in matrix-coated half, shown as brightfield and corresponding hematoxylin-and-eosin (H-E)-stained images. The full H-E image is shown in **Fig. S11**. (**B**) Comparison of CHCA matrix-coated (top row, from blue box in A) *versus* uncoated (none, bottom row, from red box in A) hippocampal horn regions excited at 488 nm and detected at various emission wavelengths ranging from 503 nm to 566 nm (from left to right), matching the blue highlighted region in the corresponding spectral data (right). (**C**) Comparison of CHCA matrix-coated (top row, from blue box in A) *versus* uncoated (none, bottom row, from red box in A) hippocampal horn regions excited at 514 nm and detected at various emission wavelengths ranging from 539 nm to 602 nm (from left to right), matching the blue highlighted region in the corresponding spectral data (right). Additional excitation wavelengths for spectral imaging are shown in **Fig. S12**. (**D**) MALDI MSI of FluoMALDI pipeline from the hippocampal horn region of the same tissue section showing MALDI images of two identified lipids, i.e., lyso-phosphatidylethanolamine (LPE) (18:2) at m/z^+^ 478.33 Da ([M+H]^+^ ion, see **Fig. S13** for tandem MS data) and PC (O-16:0) at m/z^+^ 504.35 Da ([M+Na]^+^ ion, see **Fig. S14** for tandem MS data) and their composite image acquired at 5 µm lateral spatial resolution. (**E**) MALDI MSI spectra from the two selected regions of interest (ROIs) in white boxes 1 and 2 in F showing clear intensity differences between the two regions in m/z^+^ 478.33 Da (green marker) and m/z^+^ 504.35 Da (orange marker). (**F**) Corresponding confocal laser-scanning fluorescence spectral imaging data at 488 nm excitation (left) and 514 nm excitation from the same ROIs 1 (orange line) and 2 (green line) as marked in D in the MALDI MSI data by white boxes 1 and 2.

## DISCUSSION

In summary, we have developed a new workflow, named FluoMALDI imaging, for integrated (auto)fluorescence and MALDI imaging of the same tissue section (or other biological sample), which significantly enhances (auto)fluorescence intensities through co-crystallization of endogenous or exogenous fluorophores with MALDI matrices. FluoMALDI allows for the analysis of a single tissue section across multiple modalities, which reduces the amount of sample needed and removes technical challenges associated with multimodal co-registration of adjacent tissue sections and/or independently processed samples^28, 39^. Since FluoMALDI images are collected from an identical sample prepared in an identical manner, the acquired fluorescence and MALDI MS images are inherently registered, not requiring extensive post-acquisition image registration. Since sample preparation is completed prior to any type of imaging, the FluoMALDI approach is free of any physical changes of the tissue section between fluorescence and MALDI imaging experiments, ensuring that tissue morphology remains unaltered during serial imaging steps. Therefore, combining the two imaging datasets only necessitates linear registration, which significantly simplifies alignment processes otherwise requiring complex computational methods^28, 39^.

We have shown that the FluoMALDI workflow is widely applicable to many combinations of fluorophores and matrices. The newly discovered co-crystallization phenomenon amplified the brightness of all tested fluorophore-matrix combinations in this study, up to a maximum of 79-fold. Hence, matrix deposition onto the sample allows for the enhanced detection of fluorescence signals in tissues and cells, allowing for faster fluorescence imaging with a lower threshold of detection, as demonstrated by FluoMALDI of Sharpie***®*** marker drawings, various endogenous and exogenous fluorophores, and autofluorescence from cryosections of mouse brains and kidneys. Moreover, the FluoMALDI approach resulted in greater clarity of histological features, arising from the fluorescence enhancement of endogenous compounds including FAD and PPIX, among others, in tissue sections^38, 40^. The observed fluorescence enhancement is caused by the co-crystallization process, wherein analytes, including fluorophores, are embedded within micron-sized crystals of MALDI matrices during the matrix deposition step. Both tissue fluorophores such as FAD, as well as exogenous fluorophores such as Rhodamine B/6G, could incorporate into the matrix crystals. This was evident from significant increases in fluorescence intensities in fluorophore-matrix co-crystals, as observed in real-time through time-lapse fluorescence microscopy experiments. Structural differences between pure MALDI matrix crystals and the corresponding fluorophore-matrix co-crystals suggest that the observed fluorescence enhancement is based in the co-crystallization process, which warrants deeper investigation in the future.

With FluoMALDI, we were able to collect both fluorescence and MALDI imaging data from a single tissue section in a robust and reliable manner, without compromising the imaging quality of either modality, using unmodified commercial instrumentation. Importantly, the FluoMALDI imaging approach is independent of the sequence in which consecutive fluorescence and MALDI MSI are performed, generating highly multiplexed molecular maps of mouse brain tissue with a lateral spatial resolution of down to 5 µm for both modalities, which was only limited by the spatial resolution of the MALDI MSI instrumentation available to us. With the protective matrix coating, fluorescence and MALDI imaging experiments could be conducted on different days, whereas uncoated samples will require immediate matrix deposition after fluorescence imaging to avoid degradation of tissue molecules, particularly lipids and metabolites^41^. In addition, less laser power is needed during fluorescence experiments due to the enhanced fluorescence, thereby further reducing degradation or damage to sensitive samples.

Fluorescence microscopy has long reigned as the preferred approach for multiple types of spatial biological studies, such as protein location and associations, motility, and metabolism^42-44^. Its wide-ranging applications are driven by technology advancements in confocal microscope technology^45, 46^ and multi-photon microscopy^47, 48^, allowing subcellular imaging of live and preserved samples for both targeted and untargeted studies. Recent developments in multiplexing approaches that rely on spectral imaging have further allowed for visualization and quantification of more biomarkers in complex biological systems^49, 50^. The FluoMALDI pipeline opens vast opportunities to directly integrate such powerful targeted fluorescence microscopy studies with MALDI MSI, which can detect hundreds of molecules in one experiment on one tissue section with little knowledge *a priori*. Integrating the FluoMALDI pipeline with recently reported transmission-mode MALDI-2 MSI instrumentation for subcellular resolution^9^ would be a powerful next step for future technology development, sure to drive high impact discoveries in cell biology, biomedicine, and pathology. This could be achieved by directly coupling widefield or confocal microscope optics to a MALDI MSI instrument, which would allow both imaging modalities to be executed on the same sample *in situ* or even simultaneously. This would facilitate streamlined capture of both autofluorescence or highly specific dye- or immuno-labeled fluorescence images of tissues, cells, or organelles, and MALDI images for massively multiplexed protein, metabolite, peptide, and lipid detection in the same instrument. Moreover, such combined instrumentation for the FluoMALDI pipeline would allow to directly observe, by means of fluorescence microscopy, the MALDI process in MALDI MSI.

## METHODS

### Chemicals and Reagents

Unless otherwise noted, all solvents and trifluoroacetic acid (TFA) were purchased from Sigma Aldrich (St. Louis, MO). 1,5-diaminonapthalene (DAN), α-cyano-4-hydroxycinnamic acid (CHCA), 2,5-dihydroxybenzoic acid (DHB), norharmane (nH), 9-aminoacridine (9AA) and sinapinic acid (SA) MALDI matrices were purchased from Sigma Aldrich (St. Louis, MO). The fluorophores reduced nicotinamide adenine dinucleotide (NADH) disodium, reduced NAD phosphate (NADPH) disodium, FAD disodium, Fluorescein, and Rhodamine B were obtained from Sigma Aldrich (St. Louis, MO), and Hoechst 33342, Alexa 350, Alexa 488, and Alexa 555 were purchased from Thermo Fisher (Waltham, MA). Pink Sharpie® was purchased from a local retail store. All solvents used were MS grade.

### Tissue Samples

All animal experiments were approved by the Institutional Animal Care and Use Committee (IACUC) of the Johns Hopkins University School of Medicine, which is fully accredited by the American Association for the Accreditation of Laboratory Animal care (AAALAC). Thirteen-week-old healthy control CD1 IGS mice were sacrificed. These mice were bred in the Research Animal Resources (RAR) facility at the Johns Hopkins University School of Medicine and were originally obtained from Charles River, Wilmington, DE. Following mouse sacrifice, mouse brains and kidneys were immediately harvested, frozen in liquid nitrogen vapors and stored at -80°C.

### Fluorophore Sample Preparation

A pink Sharpie® permanent marker was used to write a series of the letter “J” of approximately 5-mm by 5-mm in size onto conductive indium-tin-oxide (ITO) slides (Delta Technologies, Loveland, CO). Six common MALDI matrices were deposited onto the J’s by spraying with an HTX M5 sprayer (HTX Technologies, Chapel Hill, NC). Specifically, 50 mM of matrix at 0.05 mL/min flow rate, 10 psi, 1800 mm/min track velocity, 60°C, 3 L/min gas flow rate, 40 mm nozzle height, and 0 s drying time was sprayed to achieve a matrix density of 1.6 µg/mm^2^. We calculated matrix density according to the manufacturer’s manual: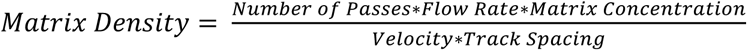 All MALDI matrix solutions were prepared in 50% acetonitrile/water + 0.1% trifluoroacetic acid. To study the effects of matrix density on fluorescence intensity, lines of pink Sharpie® of approximately 3 mm in length and 1 mm in thickness were written across ITO slides, allowed to dry under ambient conditions, and sprayed with 50 mM CHCA solution with the same spray protocols as above. The number of passes was increased to achieve densities of 0.4, 0.8, and 1.6 µg/mm^2^. Fluorophore solutions of Rhodamine B, Fluorescein, Hoechst, Alexa 350, Alexa 488, Alexa 555, NADH, NADPH, and FAD were dissolved in water at 0.2 mg/mL each. PPIX was dissolved in 50% acetonitrile/water + 0.1% trifluoroacetic acid at 0.2 mg/mL. All fluorophore solutions were spotted onto microscopy slides as 2 µl droplets and dried under ambient conditions.

### Biological Sample Preparation

Transverse (axial) mouse brain sections and coronal mouse kidney sections were cryosectioned at 10 µm thickness using a CM1860 UV Cryostat (Leica Biosystems, Wetzlar, Germany) and thaw mounted onto washed ITO slides. These sections were kept in vacuum-sealed slide mailers in the -80°C freezer until matrix deposition. The slides were allowed to warm to ambient temperature in a vacuum desiccator prior to matrix deposition using the same spray parameters as described for pink Sharpie® J’s. During tissue spraying, half of each brain or kidney section was covered by aluminum foil to remain matrix free and serve as controls. After matrix deposition, all brain and kidney slides were placed in slide mailers, wrapped with aluminum foil, vacuum-sealed and stored at -20°C until fluorescence and MALDI imaging experiments. To minimize water condensation, samples were allowed to reach room temperature prior to breaking the vacuum seal for the downstream analyses.

### Widefield Fluorescence Imaging

Images shown in **Figs. 2-3** and **S2-S4, S6** were captured using an ImageXpress Micro XLS High-Content Imager (Molecular Devices, San Jose, CA) and a Plan Apo 4x/0.2 objective. DAPI (377/50 nm excitation, 447/60 nm emission, referred to as blue fluorescence), GFP (472/30 nm excitation, 520/35 nm emission, referred to as green fluorescence), and TRITC (543/22 nm excitation, 593/40 nm emission, referred to as red fluorescence) filter sets were used to detect fluorescence images across the visible wavelength range, and a red-light emitting diode (LED) was used to capture monochromatic brightfield images. Exposure times were kept constant for a given sample type (Sharpie® images at 20 ms, tissue images at 150 ms).

### Brightfield Images

Color brightfield images in **Fig. 2** were acquired on Epson Perfection V600 Photo and in **Fig. 3A, S4, 5A** on Epson V850 Pro in 24-bit color mode at 3200 dpi or higher with backlight correction and autoexposure. Monochromatic brightfield images in **Fig. 3C, S2, S3, S6** were acquired together with the fluorescence channels using a red LED on the ImageXpress Micro XLS High-Content Imager and a Plan Apo 4x/0.2 objective.

### Polarized Light Microscopy

Images shown in **Fig. 4A** were acquired using an Olympus BX-51 upright microscope (Olympus Corporation, Tokyo, Japan) equipped with a LUCPlanFl 40x/0.6 objective and a DP-70 color CCD camera. Linear polarizers were inserted at the front focal plane of the condenser and the rear focal plane of the objective and rotated to achieve maximum extinction. This allowed birefringent materials such as the crystals of the MALDI matrix to be visualized with high contrast.

### Time-Lapse Microscopy of Co-Crystallization

Co-crystallization experiments shown in **Figs. 4B-D, S10**, and videos **V1-V4** involved mixing of equal amounts of 0.1 mM fluorophore solution with 50 mM CHCA matrix solution, both prepared in 50% acetonitrile/water + 0.1% TFA, on a glass slide. Time-lapse videos of the subsequent crystal formation and growth were acquired using an Olympus IX-81 microscope (Olympus Corporation, Tokyo, Japan) equipped with a LUCPlanFL N 20x/0.45 objective and a Hamamatsu Orca Flash 4.0 v2 sCMOS camera. A TRITC filter set was used for Rhodamine B and CHCA time-lapse acquisitions, and a GFP filter set was used for FAD time-lapse acquisitions. All imaging was performed at a rate of 10 frames per second.

### Confocal Laser-Scanning Microscopy

All confocal images in **Figs. 5, S12** were taken with a Zeiss LSM780 laser-scanning confocal microscope (Carl Zeiss AG, Oberkochen, Germany) equipped with 405, 458, 488, 514, 561, and 633 nm laser lines and a 32-channel GaAsP spectral detector. A Plan-Neofluar 10x/0.3 objective was used for all images. The instrument was configured for green emission (500-550 nm), and a tiled image of the entire brain section was acquired for both 405 and 488 nm excitation wavelengths. Next, a region of interest (ROI) around the hippocampal horn for both the matrix-coated and uncoated sides of the brain was selected for spectral imaging. Spectral images were acquired from 415-690 nm emission, in 9 nm-wide bands, using each laser line on the microscope in turn. All images were exported to ImageJ, brightness/contrast adjusted uniformly, and the “Fire” look-up table applied to better showcase intensity differences. The mean fluorescence intensity (MFI) of the entire image for each ROI was measured at each emission wavelength to generate the spectral plots, and a selection of 8 narrowband spectral images around the emission peak were used to generate the image montages. Only the 488 nm and 514 nm excitation spectral images, which produced the strongest fluorescence, are reported in **Fig.5**. All additional spectral images are shown in **Fig. S12**. For **Fig. 5F**, the same spectral measurements were taken on sub-ROIs 1 and 2 matching those used for the MALDI imaging.

### Fluorescence Image Processing and Data Analysis

Fluorescence images were exported to Fiji^51^ (ImageJ) for visualization and quantification. Tiled images were flat fielded prior to stitching to avoid grid lines. Widefield fluorescence images were also subtracted by the camera offset of 100 a.u. prior to quantification. The mean fluorescence intensity of ROIs both on and off tissue were taken to calculate the fluorescence enhancement, which is the fold change between the matrix-coated and uncoated sample acquired with the same microscope settings, after subtracting the matrix fluorescence background. Detailed methods for fluorescence image processing can be found in supporting information. **Fig. 4C** traces were generated from the raw video files in Fiji, using the following workflow: first, an 80% threshold was performed on the final timepoint to generate a mask image of fully-grown crystals. The largest 20 crystals from each video were added as ROIs, and the mean fluorescence intensity of each ROI was measured at each timepoint to generate the plot.

### MALDI Imaging Mass Spectrometry

MALDI imaging experiments in **Fig. 2-3, S4** were performed with a Bruker rapifleX MALDI TOF/TOF system (Bruker Daltonics, Bremen, Germany) equipped with a 10 kHz smartbeam 3D Nd:YAG laser at 355 nm. flexControl (version 4.0) was used to control the instrument and optimize the acquisition parameters, while flexImaging (version 5.0) was used to set up the imaging experiments and select the regions of interest. Prior to data acquisition, height adjustment (target profile generation), laser focus tuning, and external mass calibration with red phosphorus were conducted to achieve a mass error <5 PPM. For *Sharpie®* drawings in **Fig. 2**, data were acquired for m/z 100-1000 Da in reflectron positive ion mode at 50 µm raster width with 50 shots per pixel using the single laser beam scan mode, 46 µm scan range and 10 kHz frequency. The extraction voltage was set to 20 kV, lens voltage of 11.3 kV, reflectron voltages at 20.86 kV, 1.085 kV and 8.6 kV, with a delay time of 100 ns. Brain tissue MALDI images in **Fig. 3, S4** were acquired for m/z 600-1000 Da in reflectron negative ion mode at 50 µm raster width with 50 shots per pixel using the single laser beam scan mode, 46 µm scan range and 10 kHz frequency. The extraction voltage was set to -20 kV, lens voltage of -11.3 kV, reflectron voltages at -20.86 kV, -1.085 kV and -8.6 kV, with a delay time of 100 ns, while the laser fluence was optimized for each matrix. Tandem MS in **Figs. S1, S5** were conducted with the LIFT unit of the rapifleX TOF/TOF system with argon as a collision gas with an isolation window of ±1 m/z at 60% laser power boost and 4000 laser shots for fragmentation.

MALDI imaging data in **Fig. 5** were acquired on a timsTOF fleX MALDI-2 system (Bruker Daltonics, Bremen, Germany) equipped with dual ESI/MALDI source with a 10 kHz smartbeam 3D Nd:YAG laser at 355 nm with tims off. Prior to data acquisition, height adjustment (target profile generation) and laser focus tuning were conducted. Electrospray ionization (ESI) of Agilent ESI-L Tune Mix was used to perform mass calibration, followed by calibration with red phosphorus to achieve a mass error <1 PPM. MALDI parameters in qTOF mode were optimized to maximize intensity by tuning ion optics, laser intensity, and laser focus. Additional experimental values include: MALDI plate offset of 50 V, deflection 1 delta of 70 V, funnel 1 RF of 350 Vpp, funnel 2 RF 350 Vpp, multipole RF of 400 Vpp, and a collision cell energy of 10 eV, a collision RF of 1800 Vpp, quadrupole ion energy 5 eV with low mass of m/z 300, focus pre TOF transfer time of 70 μs, and a prepulse storage time of 10 μs, and both high sensitivity detection and focus mode turned on. All images were acquired for m/z 300-1300 Da in positive ion mode at 5 µm raster width with 60 shots per pixel using the single laser with beam scan disabled resulting in 5 µm field size and 10 kHz frequency. On-tissue tandem MS of m/z^+^ 478.33 Da (**Fig. S13**) and m/z^+^ 504.04 Da (**Fig. S14**) was performed on timsTOF fleX with nitrogen as a collision gas, acquiring a sum of 5 spectra using 200 shots per pixel for m/z 50-1000 Da. The laser power was set to 86% at 10 kHz frequency. The isolation window was ±1 Da, with a collision energy of 45 eV.

### MALDI MSI Data Processing and Analysis

For MALDI imaging experiments conducted on Bruker rapifleX TOF/TOF instrument shown in **Fig. 2-3**, MALDI MSI data were directly exported from flexImaging (version 5.0, Bruker Daltonics) with total ion current (TIC) normalization. For MALDI imaging experiments on timsTOF fleX shown in **Fig. 5**, MALDI MSI data were imported into SCiLS Lab (version 2023b, Bruker Daltonics) and data visualization and analysis were conducted on TIC normalized data. All spectral displays from MSI data were exported from flexImaging. Tandem MS spectra were exported from dataAnalysis (version 5.3.236, Bruker Daltonics).

### Histology

Following FluoMALDI, brain and kidney tissue sections were submerged in ethanol for 24 hours to remove MALDI matrices, followed by hematoxylin-and-eosin (H-E) staining conducted on the same tissue section using a standard protocol with Mayer’s hematoxylin solution and aqueous Eosin Y solution from Sigma Aldrich (St. Louis, MO). H-E-stained slides were imaged at 40x magnification using a NanoZoomer S210 Digital slide scanner (Hamamatsu Photonics, Shizuoka, Japan). The H-E-stained tissue sections were visualized and exported from Aperio ImageScope (version 12.4.6, Leica Biosystems, Deer Park, IL).

## ACKNOWLEDGEMENTS

The authors acknowledge the Johns Hopkins Applied Imaging Mass Spectrometry (AIMS) Core facility where all MALDI MSI experiments were performed. We also acknowledge the Johns Hopkins School of Medicine Microscopy Facility (MicFac) where all microscopy experiments were performed. We would like to thank Dr. Shruthi Shanmukha for providing the mice for this study. We further acknowledge funding from the National Institutes of Health grants R01 CA213492, R01 CA213428, R01 CA264901, and S10 OD030500.

