## Supporting Information for "FluoMALDI microscopy: matrix co-crystallization simultaneously enhances fluorescence and MALDI imaging"

**Funding information**
National Institutes of Health grants R01 CA213492, R01 CA213428, R01 CA264901, S10 OD030500.

**Table of Contents for Supporting Information**

**Supporting Methods** .......... 3
Supporting Methods: Fluorescence image processing and data analysis – additional details .......... 3
Supporting Methods: MALDI mass spectrometry imaging – additional details .......... 3
Supporting Methods: MALDI MSI data processing and analysis – additional details .......... 4

**Supporting Figures** .......... 5
Supporting Figure S1: Tandem MS spectra of Rhodamine B and pink Sharpie® .......... 5
Supporting Figure S2: FluoMALDI fluorescence enhancement of exogenous fluorophores .......... 6
Supporting Figure S3: FluoMALDI fluorescence enhancement of endogenous fluorophores .......... 7
Supporting Figure S4: FluoMALDI autofluorescence enhancement of mouse brain sections .......... 8
Supporting Figure S5: Tandem MS spectra of $m/z^-$ 885.8 Da identified as PI (38:4) .......... 9
Supporting Figure S6: FluoMALDI autofluorescence enhancement of mouse kidney sections .......... 10
Supporting Figure S7: Tandem MS spectra of $m/z^+$ 785.45 Da identified as PA (20:3_20:4) .......... 11
Supporting Figure S8: Tandem MS spectra of $m/z^-$ 621.30 Da identified as LPI (20:3) .......... 12
Supporting Figure S9: Tandem MS spectra of $m/z^-$ 906.63 Da identified as SHexCer (42:1);O3 .......... 13
Supporting Figure S10: Full length co-crystallization time course for ROD+CHCA .......... 14
Supporting Figure S11: Brightfield and H-E-stained images of brain section in Fig. 5 .......... 15
Supporting Figure S12: FluoMALDI spectral imaging for all acquired excitation wavelengths .......... 16
Supporting Figure S13: Tandem MS spectra of $m/z^+$ 478.33 Da identified as LPE (18:2) .......... 17
Supporting Figure S14: Tandem MS spectra of $m/z^+$ 504.04 Da identified as PC (O-16:0) .......... 18

**Author Contributions** .......... 19

**Supporting Methods**

**Fluorescence Image Processing and Data Analysis – Additional Details**

For fluorescence measurements, we quantified the mean pixel fluorescence intensity (MPFI), which accounted for different matrices having different fluorescence backgrounds. For quantitative measurement of Sharpie ® J markings, we measured MPFI from the entire region of the J for each matrix, solvent, and control (none). MPFI background away from the J marking was measured as well. We measured the red fluorescence intensity on and off the J markings based on these MPFI values according to the following formula: $FL\ Intensity\ (a.u.)\ = MPFI(on - J) - MPFI(off - J)$.

To calculate the fold enhancement of fluorescence intensities from the J markings, we applied the following formula: $Fold\ Enhancement = \frac{MPFI(on{-}J\ matrix)/MPFI(off{-}J\ matrix)}{MPFI(on{-}J\ none)/MPFI(off{-}J\ none)}$.

To measure fold change for different matrix densities on Sharpie ® lines, we used five randomly selected regions, placed within the lines to avoid any edge effects, to quantify MPFI. These MPFI values were used in the following formula, which is given for the example of 8 layers (8x) of CHCA matrix compared to no matrix (0x): $Fold\ Change = \frac{FL\ Intensity(8x)}{FL\ Intensity\ (0x)}$.

For quantitative measurement of fluorophore spots, we measured the green and red fluorescence intensity on and off the fluorophore spots for CHCA coated spots and uncoated controls (none). Each fluorophore spot was circled based on the brightfield images at a diameter inside the spot that avoided the dried edge effect. Quantification of all tested fluorophores with CHCA matrix was performed according to the formula: $Fold\ Change = \frac{FL\ Intensity(CHCA)}{FL\ Intensity\ (none)}$.

For quantitative measurements of brain and kidney sections, we quantified MPFI on and off tissue on both the coated and uncoated halves of brains. For on-tissue MPFI measurements, we circled the entire coated or uncoated half of the brain or kidney, or the entire uncoated brain or kidney for “none” and “solvent” controls and calculated the MPFI. For off tissue measurements, we chose five randomly selected regions off tissue to calculate MPFI. Then, we used the following formula to calculate fluorescence intensity: $FL\ Intensity\ (a.u.)\ = MPFI(on - tissue) - MPFI(off - tissue)$. The uncoated tissue halves served as internal controls and had the same values as the entirely uncoated control (none).

For calculating fold enhancement for tissue sections, the following formula was used: $Fold\ Enhancement = \frac{MPFI(on{-}tissue\ matrix)/MPFI(off{-}tissue\ matrix)}{MPFI(on{-}tissue\ none)/MPFI(off{-}tissue\ none)}$.

**MALDI Mass Spectrometry Imaging – Additional Details**

For timsTOF measurements of mouse kidney sections in **Fig. S6** in positive ion mode, prior to data acquisition, height adjustment (target profile generation), laser focus tuning, and mass calibration were conducted. Electrospray ionization (ESI) of Agilent ESI-L Tune Mix was used to perform mass calibration, followed by calibration with red phosphorus to achieve a mass error <1 PPM. MALDI parameters in qTOF mode were optimized to maximize intensity by tuning ion optics, laser intensity, and laser focus. Additional experimental values include: MALDI plate offset of 50 V, deflection 1 delta of 70 V, funnel 1 RF of 350

Vpp, funnel 2 RF 350 Vpp, multipole RF of 350 Vpp, and a collision cell energy of 10 eV, a collision RF of 2500 Vpp, quadrupole ion energy 5 eV with low mass of m/z 300 Da, focus pre TOF transfer time of 80 µs, and a prepulse storage time of 10 µs, and both high sensitivity detection and focus mode turned off. All images were acquired for m/z 300-1000 Da in positive ion mode at 50 µm raster width with 200 shots per pixel using the single laser with beam scan range of 46 µm resulting in 50 µm field size and 10 kHz frequency.

For timsTOF measurements of mouse kidney sections in **Fig. S6** in negative ion mode, prior to data acquisition, height adjustment (target profile generation), laser focus tuning, and mass calibration were conducted. ESI of Agilent ESI-L Tune Mix was used to perform mass calibration, followed by calibration with red phosphorus to achieve a mass error <1 PPM. MALDI parameters in qTOF mode were optimized to maximize intensity by tuning ion optics, laser intensity, and laser focus. Additional experimental values include: MALDI plate offset of 50 V, deflection 1 delta of -70 V, funnel 1 RF of 350 Vpp, funnel 2 RF 350 Vpp, multipole RF of 350 Vpp, and a collision cell energy of 10 eV, a collision RF of 2500 Vpp, quadrupole ion energy 5 eV with low mass of m/z 300 Da, focus pre TOF transfer time of 110 µs, and a prepulse storage time of 5 µs, and both high sensitivity detection and focus mode turned off. All images were acquired for m/z 500-1500 Da in negative ion mode at 50 µm raster width with 200 shots per pixel using the single laser with beam scan range of 46 µm resulting in 50 µm field size and 10 kHz frequency.

On-tissue tandem MS of $m/z^+$ 478.33 Da and $m/z^+$ 504.04 Da was performed on timsTOF fleX with nitrogen as a collision gas, acquiring a sum of 5 spectra using 200 shots per pixel for m/z 50-1000 Da at 10 kHz frequency. The isolation window was ±1 Da. For $m/z^-$ 906.63 Da with nH matrix, the laser power was set to 80% and the collision energy was 85 eV. For $m/z^-$ 621.30 Da with 9AA matrix, the laser power was set to 80% and the collision energy was 20 eV. For $m/z^+$ 785.45 Da with CHCA matrix, the laser power was set to 86% and the collision energy was 45 eV.

**MALDI MSI Data Processing and Analysis – Additional Details**

For MALDI imaging experiments on timsTOF fleX shown in **Fig. S6** for mouse kidney sections, MALDI MSI data were imported into SCiLS Lab (v 2023b, Bruker Daltonics) and segmentation analysis was conducted on TIC normalized data through the software's segmentation pipeline, which conducts peak selection and peak alignment and smoothing prior to segmentation using the bisecting k-means algorithm in combination with correlation distance. Segmentation maps were generated and exported from SCiLS Lab. Individual representative m/z features from each segment were identified from the segmentation analysis and the corresponding results were then visualized in flexImaging. Tandem MS spectra were exported from dataAnalysis (version 5.3.236, Bruker Daltonics).

### Supporting Figures

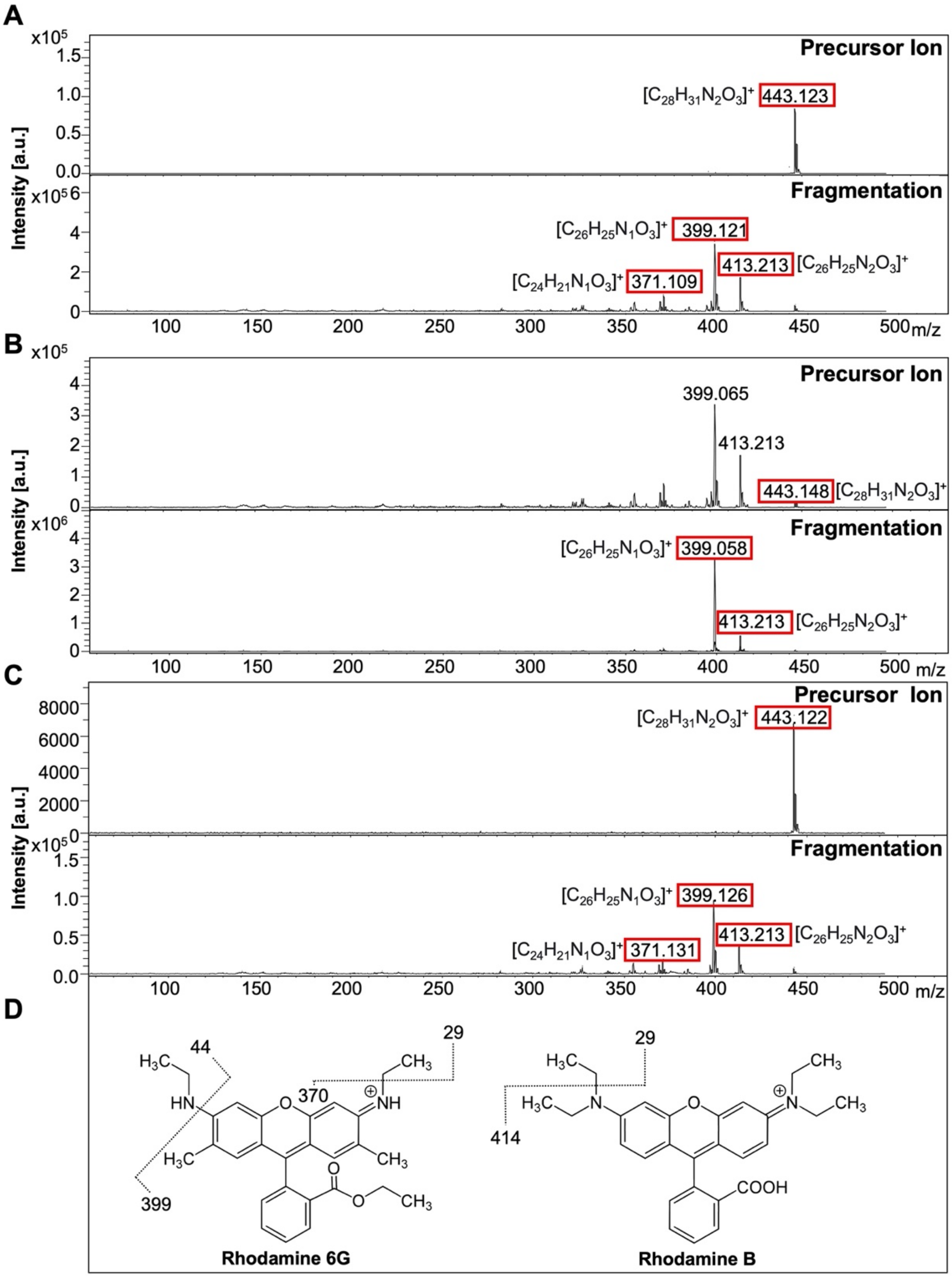


**Figure S1. Collision induced dissociation (CID) tandem MS spectra of Rhodamine B and pink Sharpie® permanent marker drawings in positive ion mode.** (**A**) Precursor ion at m/z$^+$ 443.122 Da (top) of pure Rhodamine B and resulting fragment ions (bottom). (**B**) Precursor ion at m/z$^+$ 443.122 Da (top) and fragment ions (bottom) of pink Sharpie® permanent marker drawings without matrix. Significant in source fragmentation is observed for the precursor ion. (**C**) Precursor ion at m/z$^+$ 443.122 Da (top) and fragment ions (bottom) of pink Sharpie® permanent marker drawings coated with the MALDI matrix DAN. (**D**) Chemical structures of Rhodamine 6G and B are shown, including characteristic fragmentation giving rise to diagnostic ions as boxed in red in S1A-C.

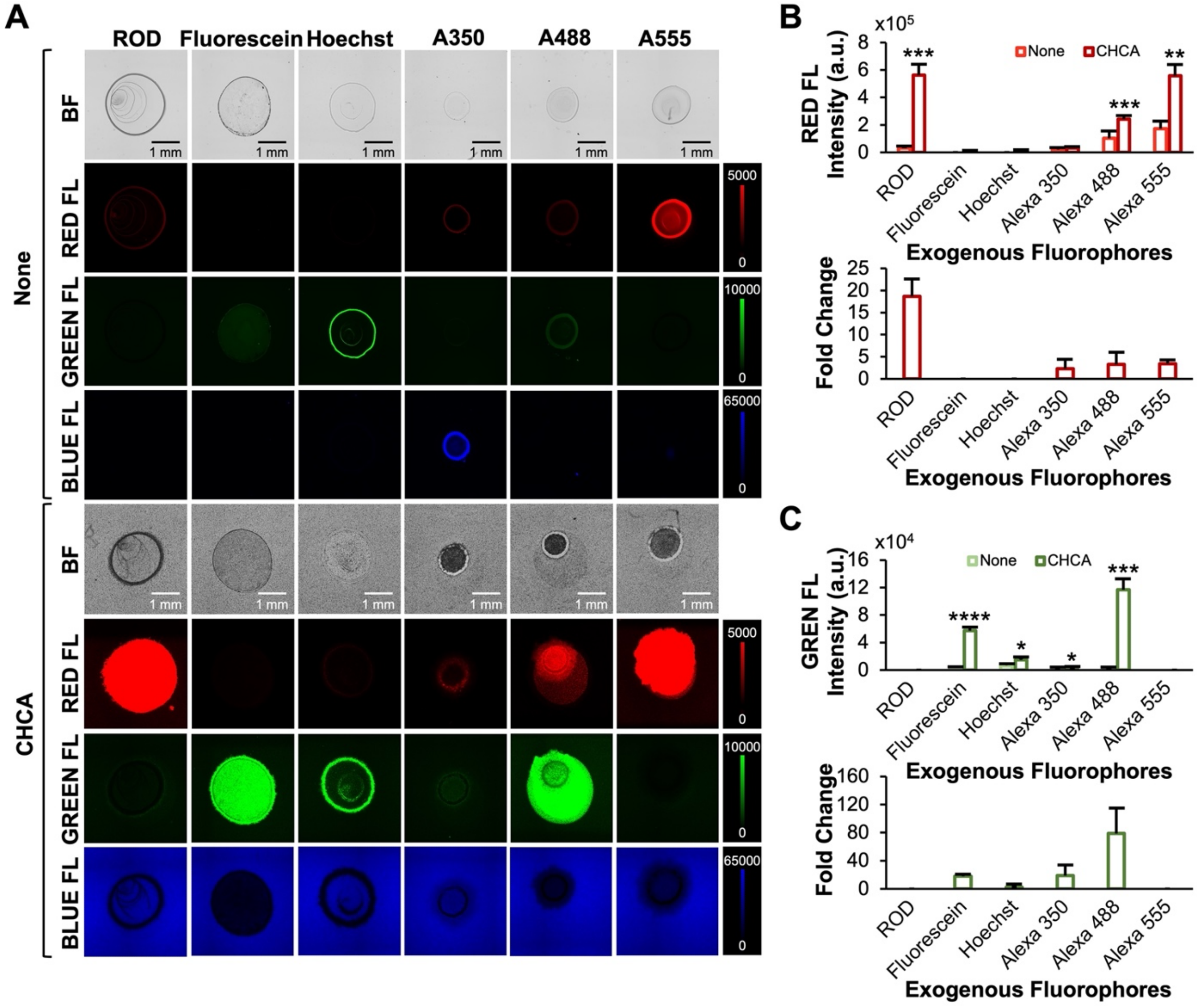


**Figure S2. Fluorescence intensity and fold change in fluorescence enhancement of exogenous fluorophores coated with CHCA matrix.** (**A**) Six common exogenous fluorophores including Rhodamine B, Fluorescein, Hoechst, Alexa Fluor 350 (A350), Alexa Fluor 488 (A488), and Alexa Fluor 555 (A555) were spotted onto slides at equal concentrations, dried, and sprayed with CHCA matrix at 1.6 μg/mm$^2$ density. Red, green, and blue epifluorescence images were acquired from uncoated fluorophores (none, top) and CHCA-coated fluorophores (bottom). (**B**) Quantification of red fluorescence intensities and fold change from CHCA-coated exogenous fluorophores compared to respective uncoated controls, n=3. (**C**) Quantification of green fluorescence intensities and fold change from CHCA-coated exogenous fluorophores compared to respective uncoated controls, n=3. Blue fluorescence intensities were not quantified because of high innate blue fluorescence of CHCA matrix causing high background signal. All quantitative data are shown as mean values ± standard error of three independent experiments. * $p<0.05$, ** $p<0.01$, *** $p<0.001$. Abbreviations: BF, brightfield; FL, fluorescence; ROD, Rhodamine B.

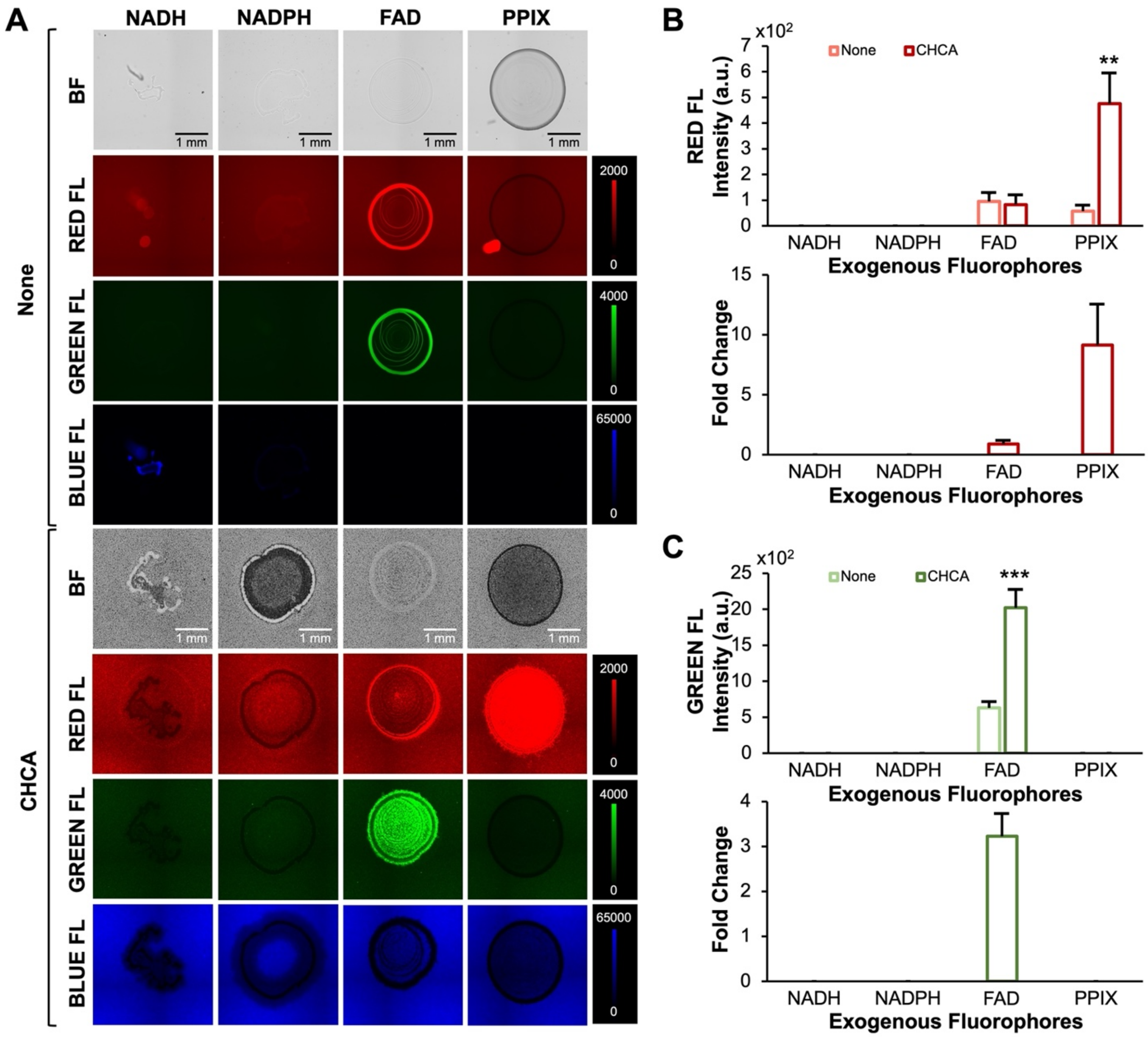


**Figure S3. Fluorescence intensity and fold change in fluorescence enhancement of endogenous fluorophores coated with CHCA matrix.** (**A**) Four common endogenous fluorophores including reduced nicotinamide adenine dinucleotide (NADH), reduced nicotinamide adenine dinucleotide phosphate (NADPH), flavin adenine dinucleotide (FAD), and protoporphyrin IX (PPIX) were spotted onto slides at equal concentrations, dried, and sprayed with CHCA matrix at 1.6 μg/mm$^2$ density. Red, green, and blue epifluorescence images were acquired from uncoated fluorophores (none, top) and CHCA-coated fluorophores (bottom). (**B**) Quantification of red fluorescence intensities and fold change from CHCA-coated endogenous fluorophores compared to respective uncoated controls, n=3. (**C**) Quantification of green fluorescence intensities and fold change from CHCA-coated endogenous fluorophores compared to respective uncoated controls, n=3. Blue fluorescence intensities were not quantified because of high innate blue fluorescence of CHCA matrix causing high background signal. All quantitative data are shown as mean values ± standard error of three independent experiments. ** p<0.01, *** p<0.001. Abbreviations: BF, brightfield; FL, fluorescence.

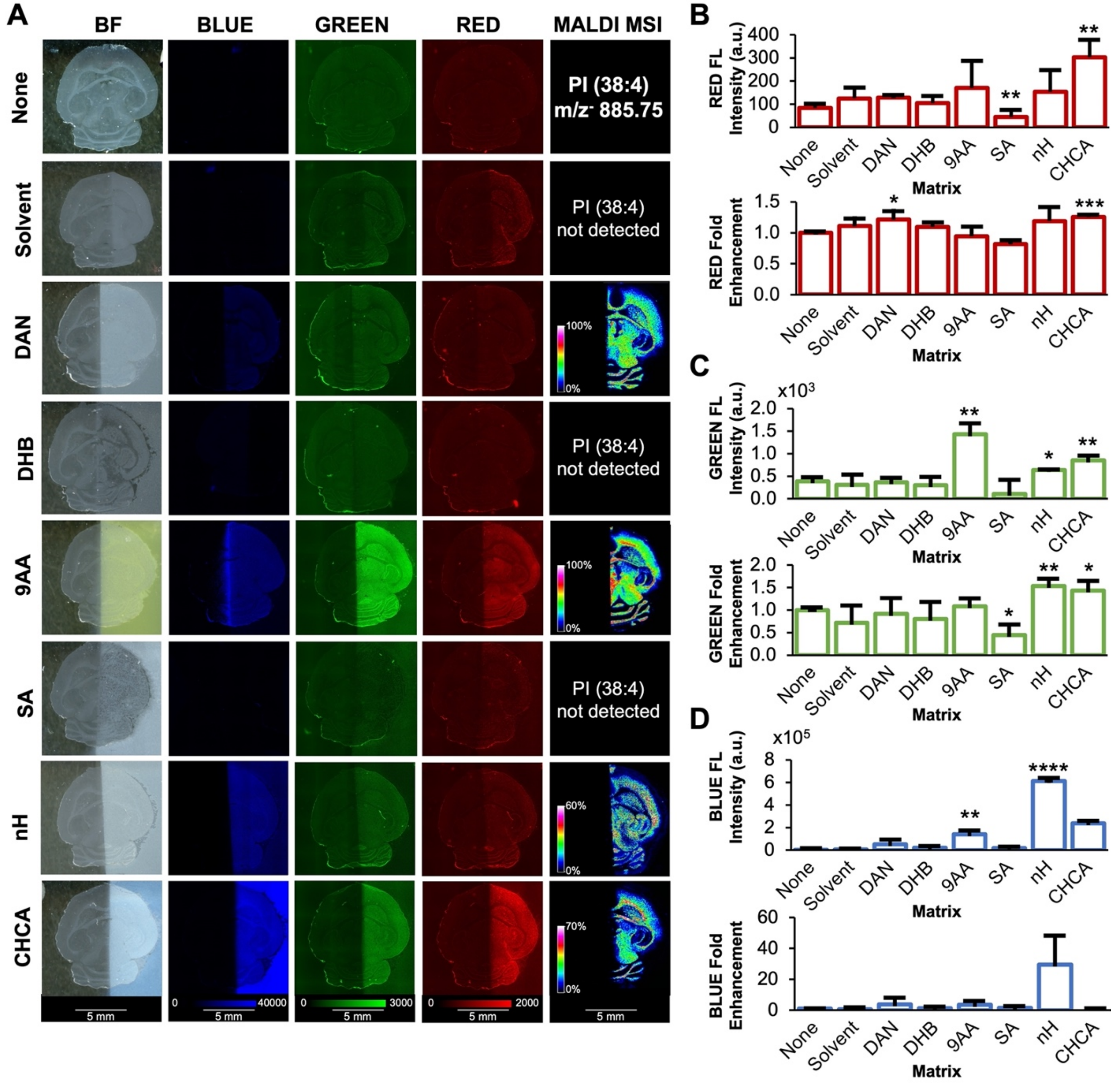


**Figure S4. MALDI matrix coating of mouse brain tissue sections increases tissue autofluorescence intensity.** (**A**) Imaging results of transverse (axial) mouse brain tissue sections showing, in columns from left to right, brightfield, green fluorescence, red fluorescence, and MALDI imaging in negative ion mode displaying $m/z^-$ 885.75 Da, which was identified by tandem MS as phosphatidylinositol (PI) (38:4) ($[M\text{-}H]^-$, see **Fig. S5** for tandem MS data). Rows show uncoated controls and coating with various MALDI matrices at 1.6 μg/mm$^2$ density. Fluorescence and MALDI images are shown on the same intensity scale per column. Quantification of (**B**) red, (**C**) green, and (**D**) blue fluorescence intensities from (A), n=3. All quantitative data are shown as mean values ± standard error of three independent experiments. * $p<0.05$, ** $p<0.01$, *** $p<0.001$. Abbreviations: BF, brightfield; FL, fluorescence.

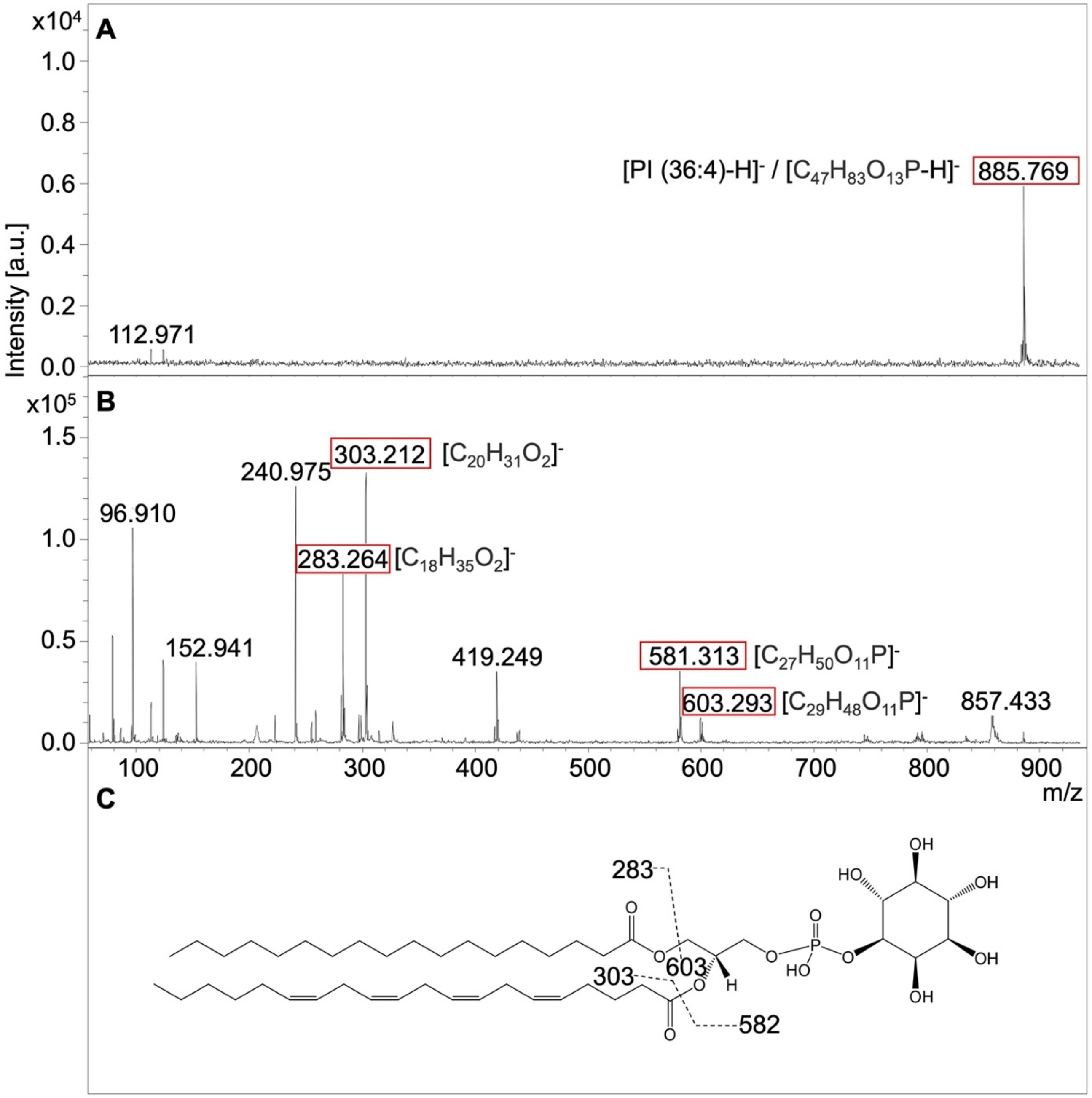


**Figure S5. Negative ion mode tandem MS spectra of m/z⁻ 885.8 Da identified as PI (38:4), [M-H]⁻.** (**A**) Precursor ion at m/z⁻ 885.8 Da, and (**B**) fragmentation of m/z⁻ 885.8 Da. Chemical structures of characteristic fragments (boxed in red) are shown.

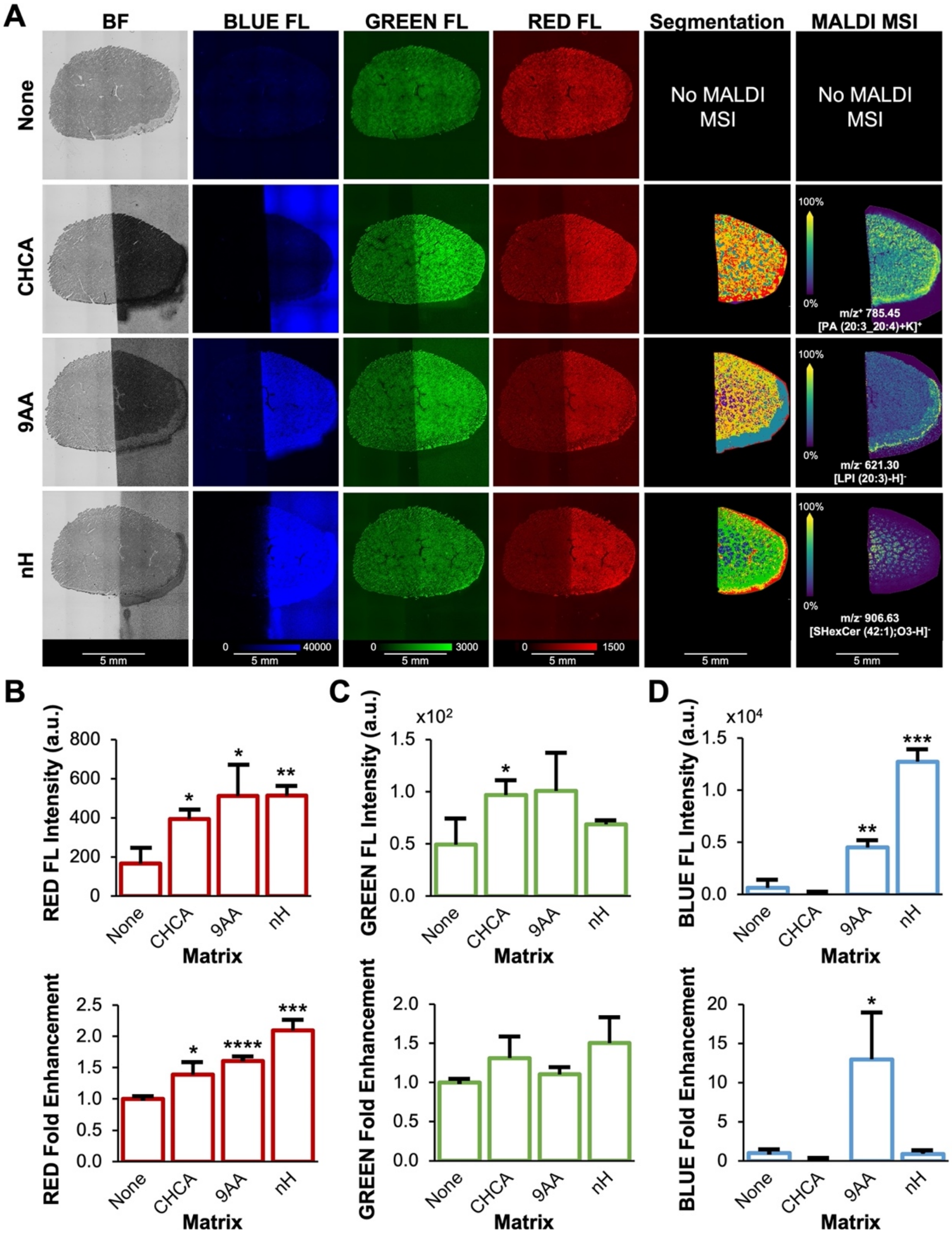


**Figure S6. MALDI matrix coating of mouse kidney tissue sections increases tissue autofluorescence intensity.** (**A**) Imaging results of coronal mouse kidney tissue sections showing, in columns from left to right, brightfield, blue fluorescence, green fluorescence, red fluorescence, segmentation analysis from MALDI imaging, MALDI imaging displaying $m/z^+$ 785.45 Da for CHCA matrix identified by on-tissue tandem MS as phosphatidic acid (PA) (20:3_20:4) $[M+K]^+$, (see **Fig. S7** for tandem MS data), $m/z^-$ 621.30 Da for 9AA matrix identified by on-tissue tandem MS as lyso-phosphatidylinositol (LPI) (20:3) $[M-H]^-$ (see **Fig. S8** for tandem MS data), and $m/z^-$ 906.63 Da for nH matrix identified by on-tissue tandem MS as sulfatide (SHexCer) (42:1);O3 $[M-H]^-$ (see **Fig. S9** for tandem MS data). Rows show uncoated control and coating with various MALDI matrices at 1.6 µg/mm$^2$ density. Fluorescence and MALDI images are shown on the same intensity scale per column. Quantification of (**B**) blue, (**C**) green, and (**D**) red fluorescence intensities from (A), n=3. All quantitative data are shown as mean values ± standard error of three independent experiments. * $p<0.05$, ** $p<0.01$, *** $p<0.001$, **** $p<0.0001$. Abbreviations: BF, brightfield; FL, fluorescence.

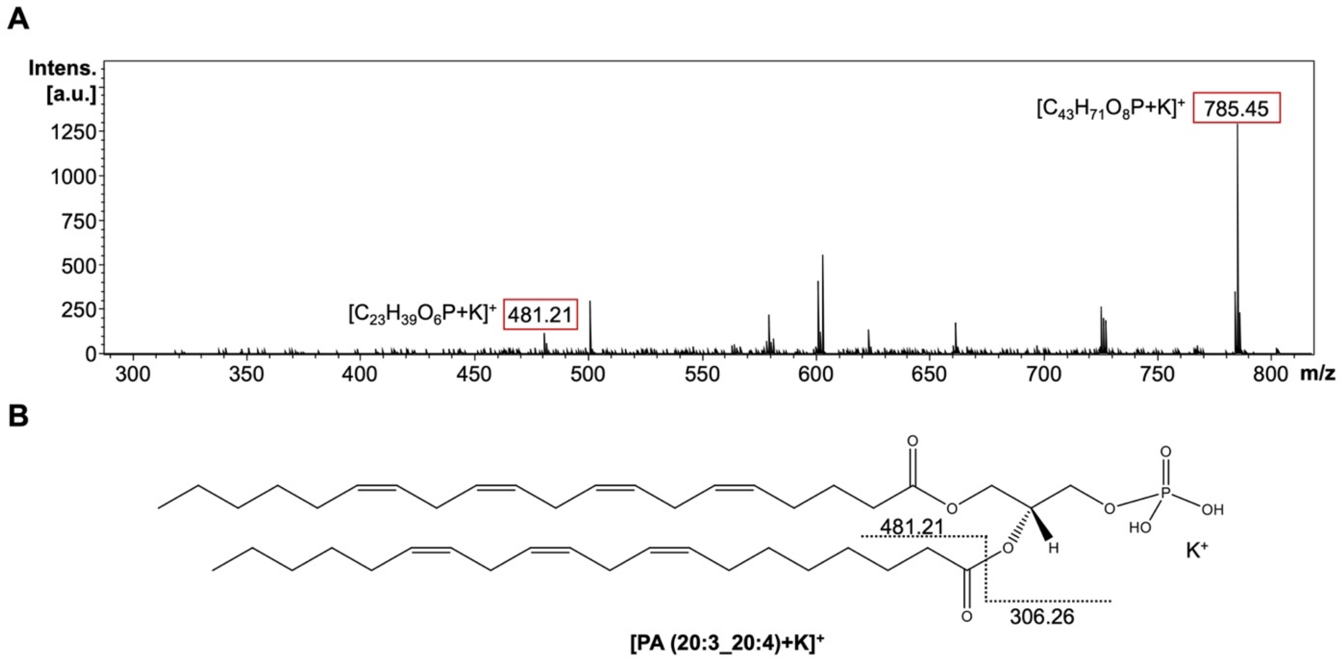


**Figure S7. Positive ion mode tandem MS spectra of m/z$^+$ 785.45 Da identified as PA (20:3_20:4), [M+K]$^+$.**
(**A**) Precursor ion and fragmentation of m/z$^+$ 785.45 Da. (**B**) Chemical structure and fragmentation are shown based on characteristic fragments (boxed in red).

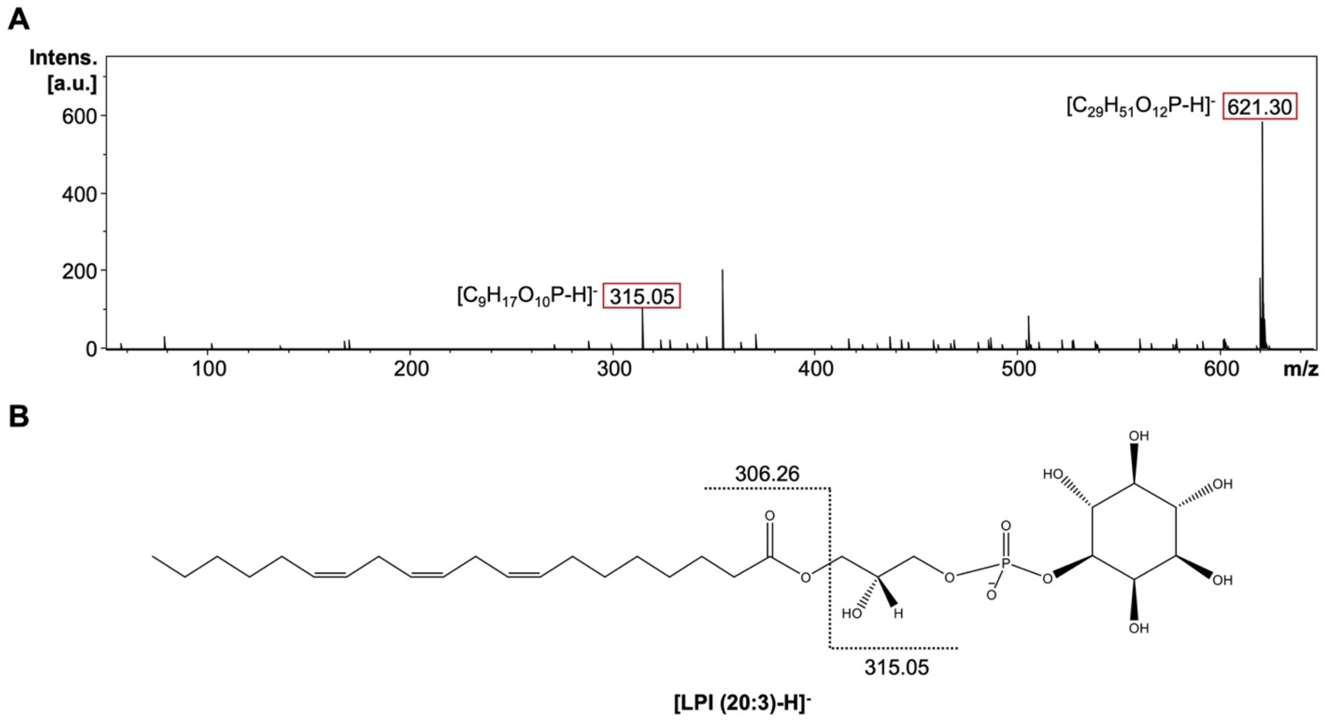


**Figure S8. Negative ion mode tandem MS spectra of m/z$^-$ 621.30 Da identified as LPI (20:3), [M-H]$^-$.**
(**A**) Precursor ion and fragmentation of m/z$^-$ 621.30 Da. (**B**) Chemical structure and fragmentation are shown based on characteristic fragments (boxed in red).

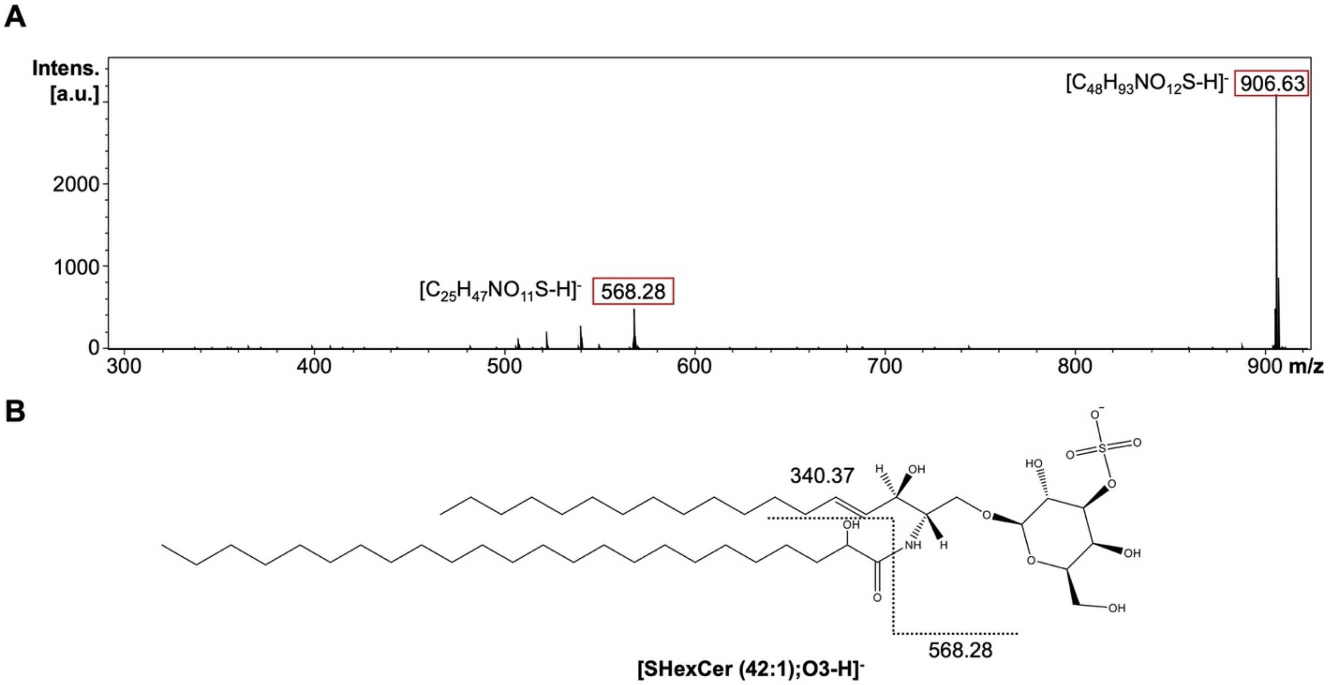


**Figure S9. Negative ion mode tandem MS spectra of m/z⁻ 906.63 Da identified as SHexCer (42:1);O3, [M-H]⁻.**
(**A**) Precursor ion and fragmentation of m/z⁻ 906.63 Da. (**B**) Chemical structure and fragmentation are shown based on characteristic fragments (boxed in red).

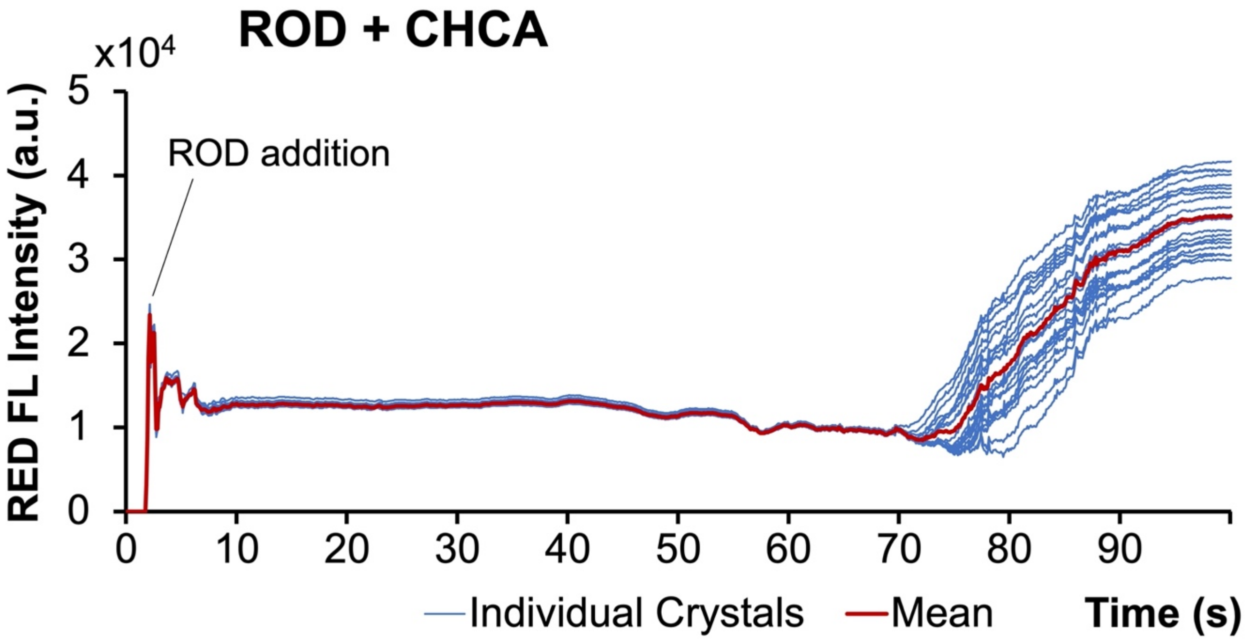


**Figure S10. Co-crystallization of fluorophores with matrices – full length time course for ROD+CHCA.** Full length time course of fluorescence intensity quantification of the 20 largest crystals for (co-)crystallization of ROD+CHCA. Blue lines represent individual crystals' fluorescence intensities, red lines are mean crystal fluorescence intensities. Time point of adding ROD to matrix solution on microscope slide is pointed out. This data is also available as **video V4**.

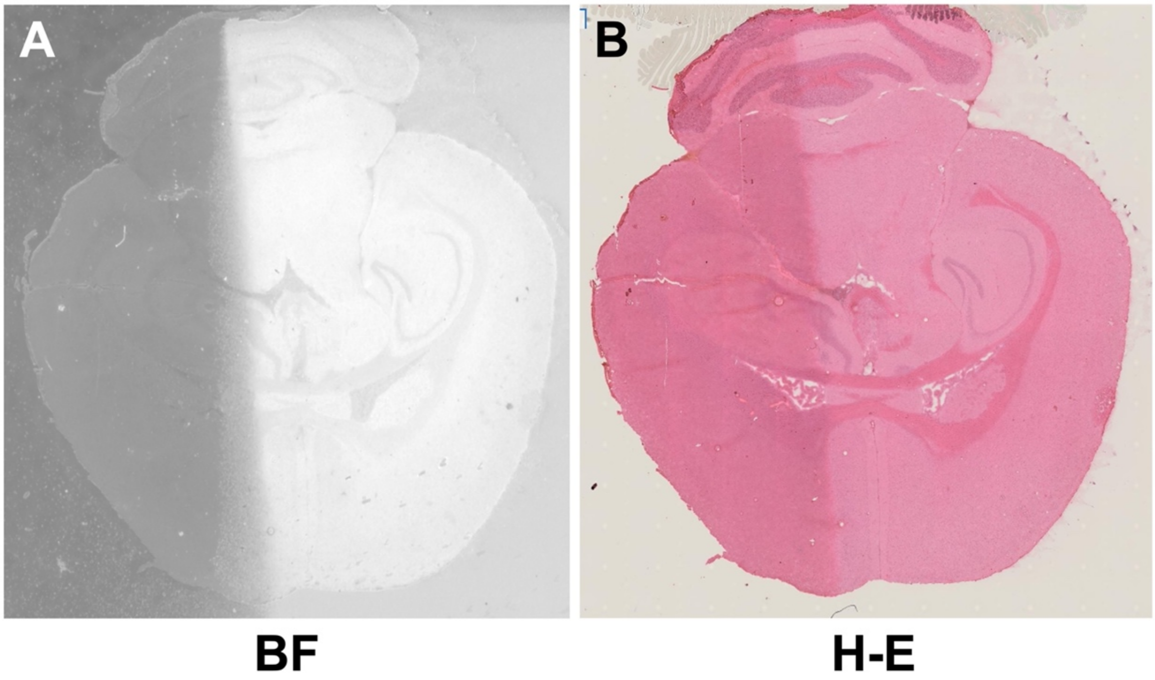


**Figure S11. Brightfield and H-E-stained images of transverse (axial) half-coated brain section shown in Fig. 5.** (**A**) Brightfield and (**B**) H-E-stained images of full brain section utilized in confocal experiments, corresponding to the data shown in **Fig. 5**.

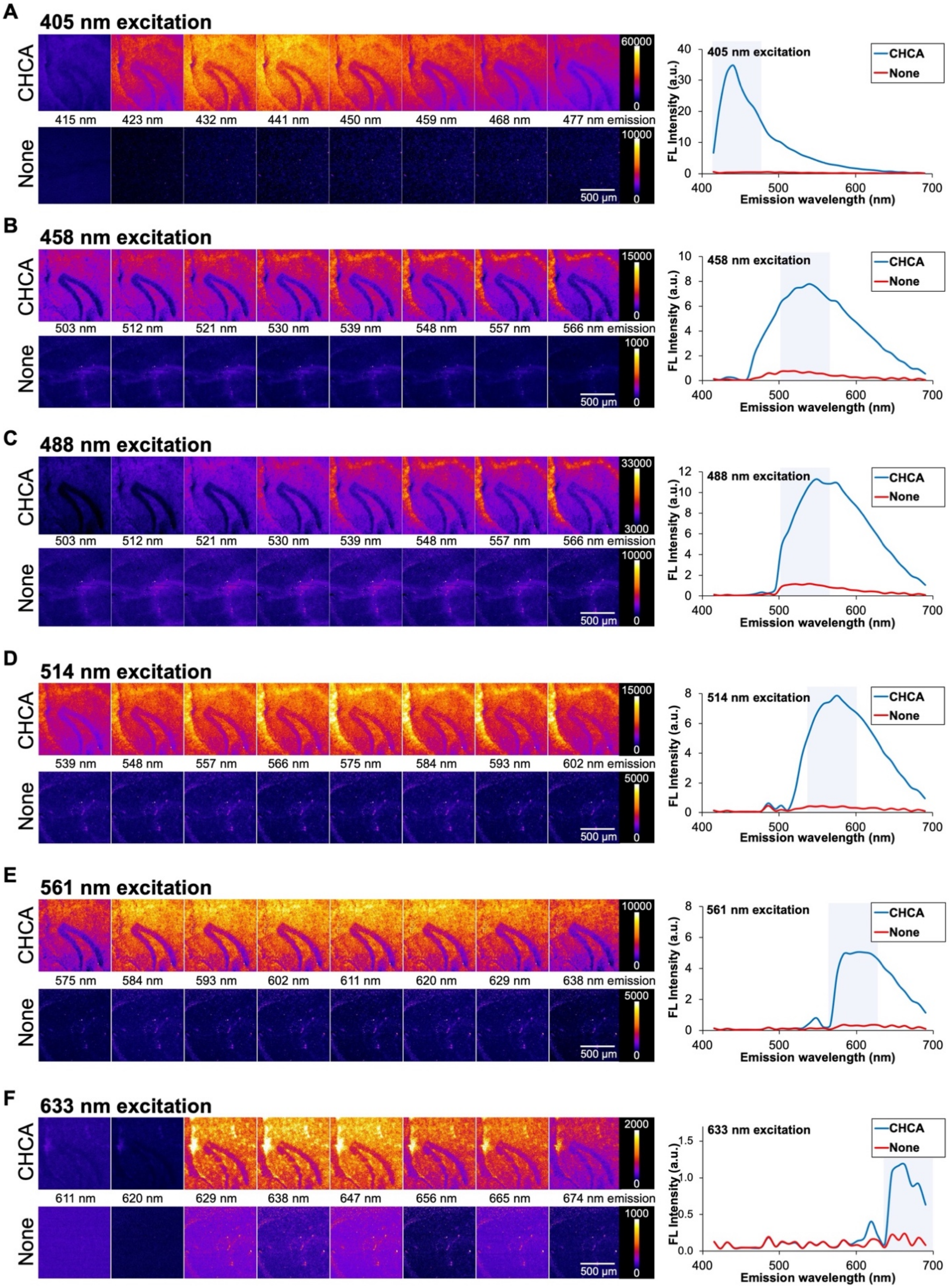


**Figure S12. FluoMALDI pipeline using spectral imaging with laser-scanning confocal microscopy and MALDI MSI on mouse brain tissue sections half-coated with CHCA**. Comparison of CHCA matrix-coated (top row, from blue box in **Fig. 5A**) *versus* uncoated (none, bottom row, from red box in **Fig. 5A**) hippocampal horn regions excited at (**A**) 405 nm, (**B**) 458 nm, (**C**) 488 nm, (**D**) 514 nm, (**E**) 561 nm, (**F**) 633 nm, and detected at various emission wavelengths ranging from 415 nm to 690 nm (from left to right), matching the blue highlighted regions in the corresponding spectral data (right).

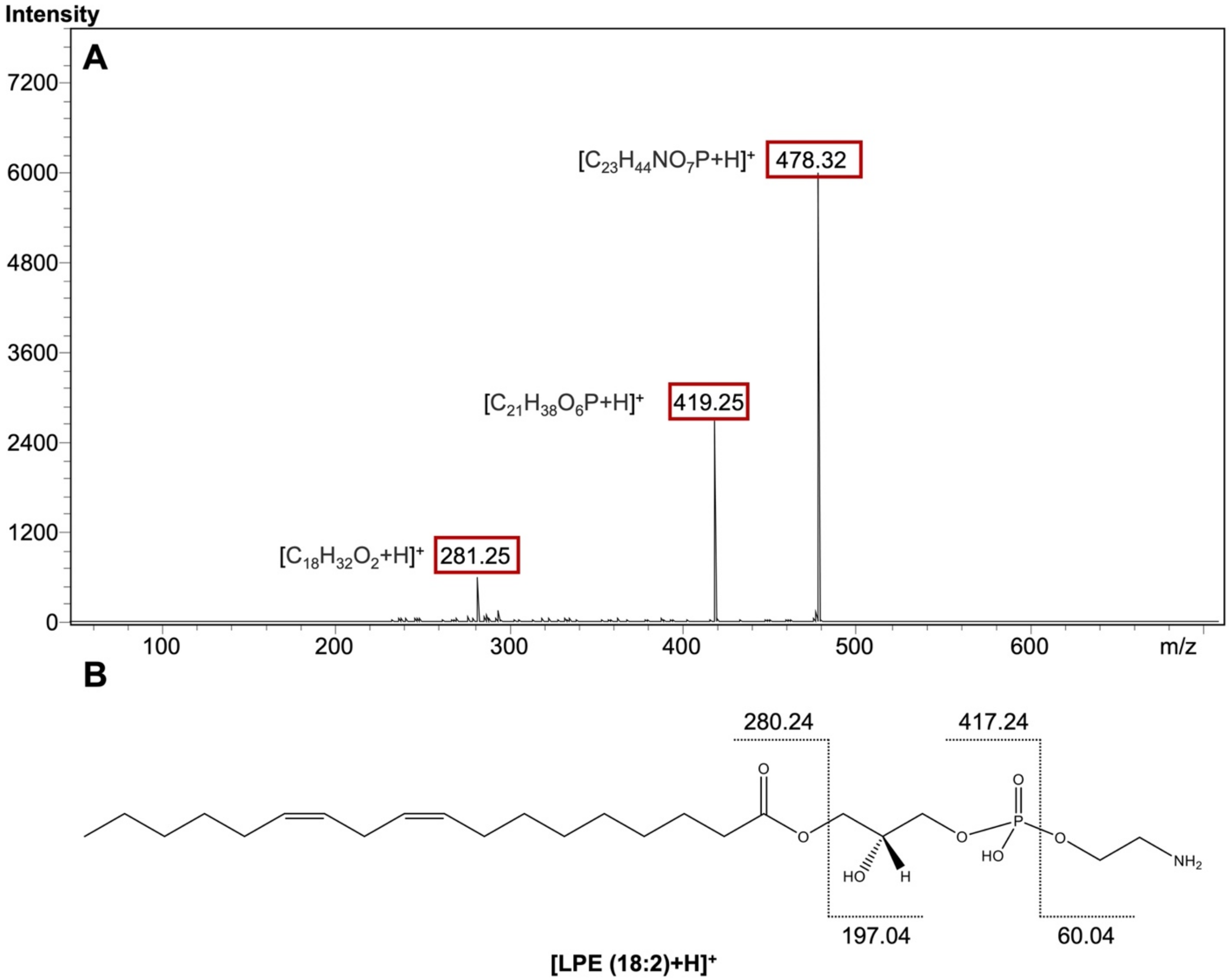


**Figure S13. Positive ion mode tandem MS spectra of m/z⁺ 478.33 Da identified as LPE (18:2), [M+H]⁺.**
(**A**) Precursor ion and fragmentation of m/z⁺ 478.33 Da. (**B**) Chemical structure and fragmentation are shown based on characteristic fragments (boxed in red).

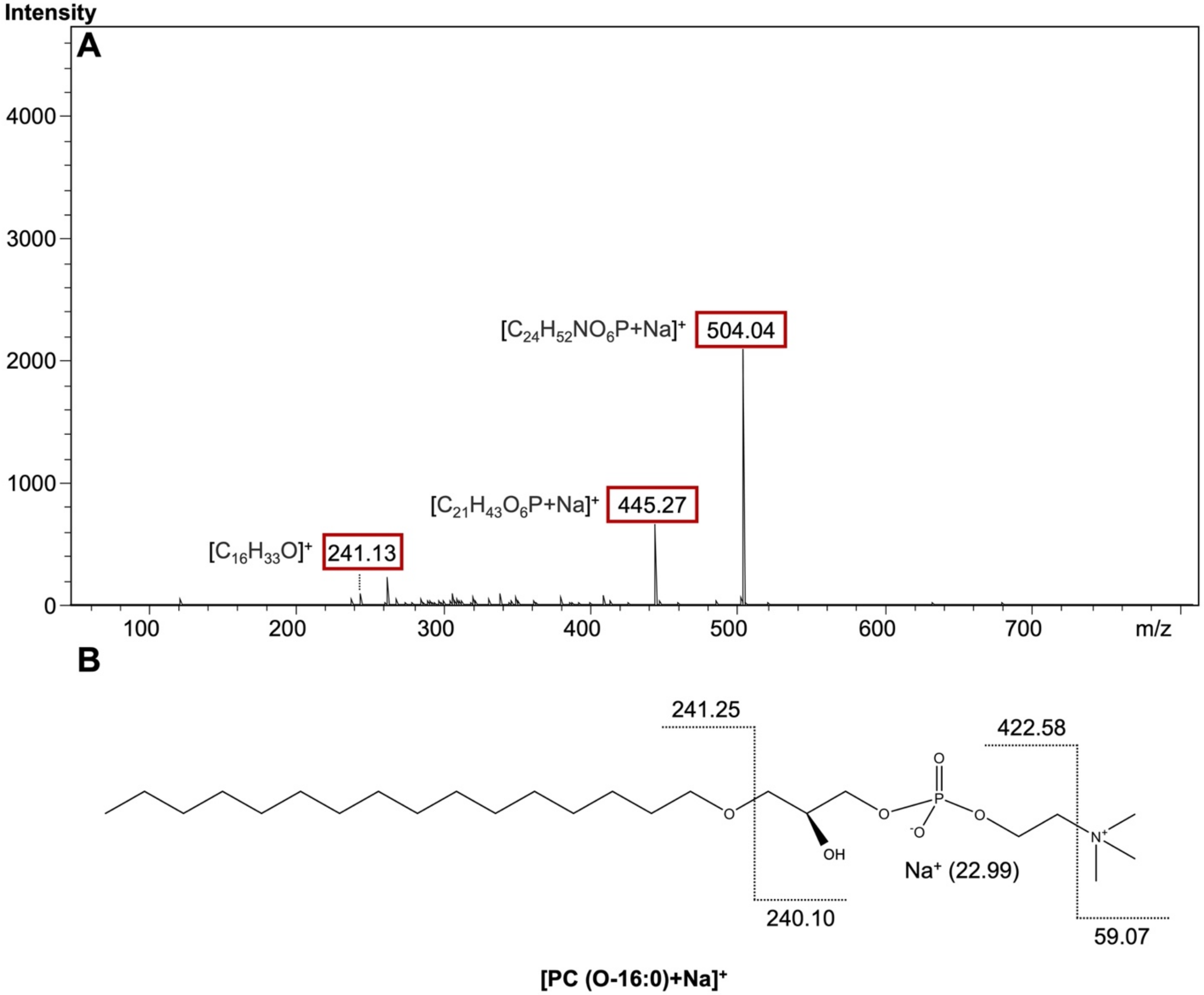


**Figure S14. Positive ion mode tandem MS spectra of m/z⁺ 504.04 Da identified as PC (O-16:0), [M+Na]⁺.**
(**A**) Precursor ion and fragmentation of m/z⁺ 504.04 Da. (**B**) Chemical structure and fragmentation are shown based on characteristic fragments (boxed in red).

### Author Contributions

KG initiated, directed, and supervised the study. KG, SK, HWF, EY, and IB conceptualized the study. EY and XES carried out sample preparation. HWF, XES and LAR performed fluorescence microscopy experiments, data processing, and analysis. EY, CMT, XES, DRB, and CCJ carried out MALDI MSI experiments, data processing, and analysis. EY, XES, DRB, CCJ, and CMT performed H-E staining and slide-scanning. EY, XES, JHK, IB, SK, HWF, and KG designed the figures. EY, XES, and KG wrote the manuscript with support from HWF. All authors contributed to, provided critical feedback, reviewed, and approved the final manuscript.
